# Characterization of the glycoproteins of novel fish influenza B-like viruses

**DOI:** 10.1101/2025.05.08.652883

**Authors:** Gagandeep Singh, Jiachen Huang, Disha Bhavsar, Kirill Vasilev, James A. Ferguson, Geert-Jan Boons, Viviana Simon, Robert P. de Vries, Julianna Han, Andrew Ward, Florian Krammer

## Abstract

Novel influenza-like virus sequences previously identified in fish and amphibians were found to cluster as a sister clade of influenza B viruses, but have thus far remained uncharacterized. We demonstrate that salamander influenza-like virus (SILV) HA is functionally divergent from influenza B virus HA and does not bind to α 2,3- and α2,6-linked sialic acids. However, the HAs of Siamese algae-eater influenza-like virus (SAEILV) and chum salmon influenza-like virus (CSILV) bind to α2,3 linked sialic acid. Furthermore, SAEILV HA binds to sialyated Lewis X, is activated by human airway enzymes and is fusogenic at a wide range of pH conditions. SAEILV NA has a highly conserved active site and a similar structure to other known NAs. We also determined the cryo-electron microscopy structure of the HA of a previously described virus from the same sister clade, the Wuhan spiny eel influenza virus (WSEIV). Importantly, no cross-reactive antibodies against these HAs or NAs were found in the human serum, suggesting that humans are immunologically naïve to these viruses.

**One sentence summary:** Novel influenza like-viruses, displayed different target receptor specificity and limited antigenic conservation of the HA and NA relative to influenza B Viruses.

## Introduction

Influenza viruses are highly contagious respiratory pathogens responsible for seasonal epidemics and occasionally, global pandemics, posing a significant threat to public health and the economy worldwide. Influenza viruses are classified into four distinct types: A, B, C, and D(*1, 2*). Waterbirds and shorebirds serve as natural reservoirs for influenza A viruses, though zoonotic spillover events have resulted in influenza A virus transmission to a wide variety of avian and mammalian species, including humans, terrestrial birds, pigs, poultry, horses, cats, cows, bats, marine mammals etc.(*3, 4*). Influenza A H1N1 and H3N2 virus strains are currently circulating in humans but 19 HA subtypes and 11 NA subtypes have been detected in the animal reservoir. The introduction of other subtypes - to which humans have little to no immunity- into the population carries the potential to cause devastating pandemics(*1*). In contrast, the influenza B virus is relatively stable due to its limited antigenic drift and narrow host range(*5–7*). Influenza B viruses are primarily restricted to human hosts and are known to cause seasonal and epidemic outbreaks of influenza(*8*). However, sporadic reports of infections detected in harbor and grey seals as well as bamboo rats have raised questions about the possibility of non-human influenza B virus reservoirs(*5, 9–11*). Phylogenetic analysis of viruses isolated from seals suggests that the influenza B virus detected there was of human origin(*12*). Thus far, there has been limited evidence showing sustained animal reservoirs of influenza B virus strains and there have been no documented cases of influenza B virus transmission from animals to humans.

Despite extensive research on the characterization of influenza viruses, our overall understanding of the true diversity and evolutionary history of vertebrate RNA viruses, particularly concerning those found outside of mammalian and avian hosts, remains limited. A 2018 study identified 214 new vertebrate-associated RNA viruses across more than 186 host species by employing a large-scale meta-transcriptomic approach. Novel influenza-like viruses discovered in fish and amphibians included Wuhan Asiatic toad, Wuhan spiny eel and Wenling hagfish influenza-like viruses(*13*) of which the Wuhan spiny eel virus (WSEIV) appeared to be influenza B virus-like(*14*).

Recently, Parry *et al.* identified the complete coding segments of five divergent vertebrate influenza-like viruses. Two amphibian influenza-like viruses exhibit relatively high pairwise amino acid identity to segments from the influenza D virus. The other three viruses were found to be closely related to influenza B virus: Salamander influenza-like virus (SILV), Siamese algae-eater influenza-like virus (SAEILV), and chum salmon influenza-like virus (CSILV), detected in Mexican walking fish (*Ambystoma mexicanum*) and plateau tiger salamander (*Ambystoma velasci*), Siamese algae-eater fish (*Gyrinocheilus aymonieri*), and chum salmon (*Oncorhynchus keta)*, respectively. Furthermore, the genome arrangements of these three viruses are similar to those of influenza A and B viruses, containing eight segments (PB1, PB2, PA, HA, NP, NA, M, and NS). Sequence alignment and phylogenetic analysis of the coding regions indicate that these three viral genomes display the closest similarities to the influenza B virus species. The amino acid percentage identity between SILV and influenza B viruses ranges from 75.57% (PB1) to 26.66% (NS), between SAEILV and influenza B viruses it ranges from 69.46% (PB1) to 24.56% (NS), and between CSILV and influenza B viruses it ranges from 69.50% (PB1) to 34.53% (M). The surface glycoproteins, hemagglutinin (HA), and the neuraminidase (NA) of SILV exhibit 43% to 46% identity, while those of SAEILV show 34% to 43% identity, and CSILV HA and NA display 32% to 38% amino acid identity to respective HA and NA of influenza B viruses(*15*). These findings suggest the existence of a vast number of uncharacterized influenza B-like viruses, and highlight our limited understanding of the origin, evolution and cross-species transmission of these viruses.

The viral surface glycoproteins HA and NA play a critical role in the life cycle of the influenza A and B virus. HA mediates viral attachment and entry into host cells through interactions with sialic glycans on the cell surface(*16, 17*). NA cleaves the glycosidic linkage between the terminal sialic acids and galactose, facilitating the release of viral progeny from the cell surface(*18*). Influenza B virus HA can interact with both α2,3- and α2,6-linked *N*-acetylneuraminic acid (Neu5Ac) (*19*). In contrast, WSEIV, an influenza B-like virus, lacks the ability to bind to the canonical influenza B virus receptors. Instead, WSEIV utilizes a sialylated ganglioside, GM2, as its receptor(*14*). Based on NA sequence comparisons, the nine subtypes of the influenza A virus NAs can be categorized into two groups: group 1 (N1, N4, N5, N8) and group 2 (N2, N3, N6, N7, N9). Interestingly, the NA-like proteins from bat influenza viruses H17N10 and H18N11 do not exhibit NA activity despite their structural similarity with authentic NAs(*20–22*). However, the NA protein of, WSEIV, has NA activity similar to the influenza B virus NA, despite low sequence identity(*14, 23*). Collectively, these findings underscore the need for studies characterizing novel influenza B-like viruses in under-sampled hosts, as well as for understanding the functionalities and antigenic properties of these viruses. To better understand the complexity and divergence of HAs and NAs from genetically distinct influenza B-like viruses, we report the functional, antigenic, and structural characterization of the salamander, Siamese algae-eater, and chum salmon influenza B-like virus glycoproteins and structural characterization of the HA of the previously described Wuhan spiny eel influenza virus.

## Results

### Phylogenetic and comparative sequence analysis

The phylogenetic analysis of representative influenza A and B virus HA and NA genetic sequences with the HA and NA of SILV, SAEILV, CSILV, and WSEIV indicated that these novel influenza-like viruses were more closely related to influenza B viruses than influenza A viruses (Fig.1A, B). The position of these sequences strongly suggests that these viruses have a long evolutionary history in vertebrates. Identical amino acid residues of each glycoprotein revealed by paired sequence analysis with the reference strain B/Brisbane/60/2008 (B/Bris/08), were highlighted in blue on top of the structures of B/Bris/08 HA and NA to indicate conserved patches. Compared to the HA and NA of B/Bris/08, SILV showed ∼42% (HA) and ∼44% (NA) amino acid identity (Fig. 1C, F), SAEILV was ∼30% (HA) and ∼40% (NA) identical (Fig. 1D, G), and CSILV ∼28% (HA) and ∼34% (NA) identical (Fig.1E, H). Multiple sequence alignment of the novel influenza-like viruses with the influenza B/Malaysia/2506/2004 virus HA and NA showed distinct glycosylation patterns and receptor binding site (RBS) residues for each strain (Fig. 1I). In contrast, comparison of the NA sequences demonstrated that residues within the enzymatic active site are highly conserved among influenza B-like viruses, along with the regions in the immediate vicinity of the enzymatic site (Fig 1 F-H, 1J). In the context of the HA, we noticed a lack of conservation in the residues that form the sialic acid-interacting RBS(*24, 25*), suggesting a potentially altered receptor binding profile for these influenza B-like HAs. Furthermore, these HA sequences appeared to have a reduced number of putative N-linked glycosylation sites compared to the B/Malaysia/2506/2004 HA, while the N336 glycan remained conserved in all the sequences analyzed. To analyze the extend of glycosylation, the HAs were expressed as recombinant protein trimers in insect cells using the baculovirus expression system as described in prior studies(*26*) and analyzed with and without PNGaseF treatment via sodium dodecyl sulfate– polyacrylamide gel electrophoresis (SDS-PAGE) in which an increased electrophoretic mobility indicates a loss of molecular weight due to the removal of N-linked glycans. As a control we used recombinant B/Malaysia/2506/2004 HA, which is part of the B/Victoria/2/1987-like lineage, also expressed using the baculovirus expression system as described previously(*27*). Compared to SILV, SAEILV and CSILV HAs, B/Malaysia/2506/2004 HA showed a greater change in molecular weight after deglycosylation (Fig. 1K). This confirms the sequence-based observation that fewer glycosylation sites are present on these HAs. From an antigenic perspective, the target epitope of the pan-influenza virus HA monoclonal antibody (mAb) CR9114 also exhibited some mismatches(*25*). However, a substantial number of matched residues were found in the stalk domain of the HA, consistent with previous findings of higher levels of conservation in this region across all influenza virus HAs(*28*). Significant mismatches were observed in the region immediately upstream of the fusion peptide, which is largely conserved and encompasses the proteolytic cleavage site essential for HA activation.

**Fig. 1.**
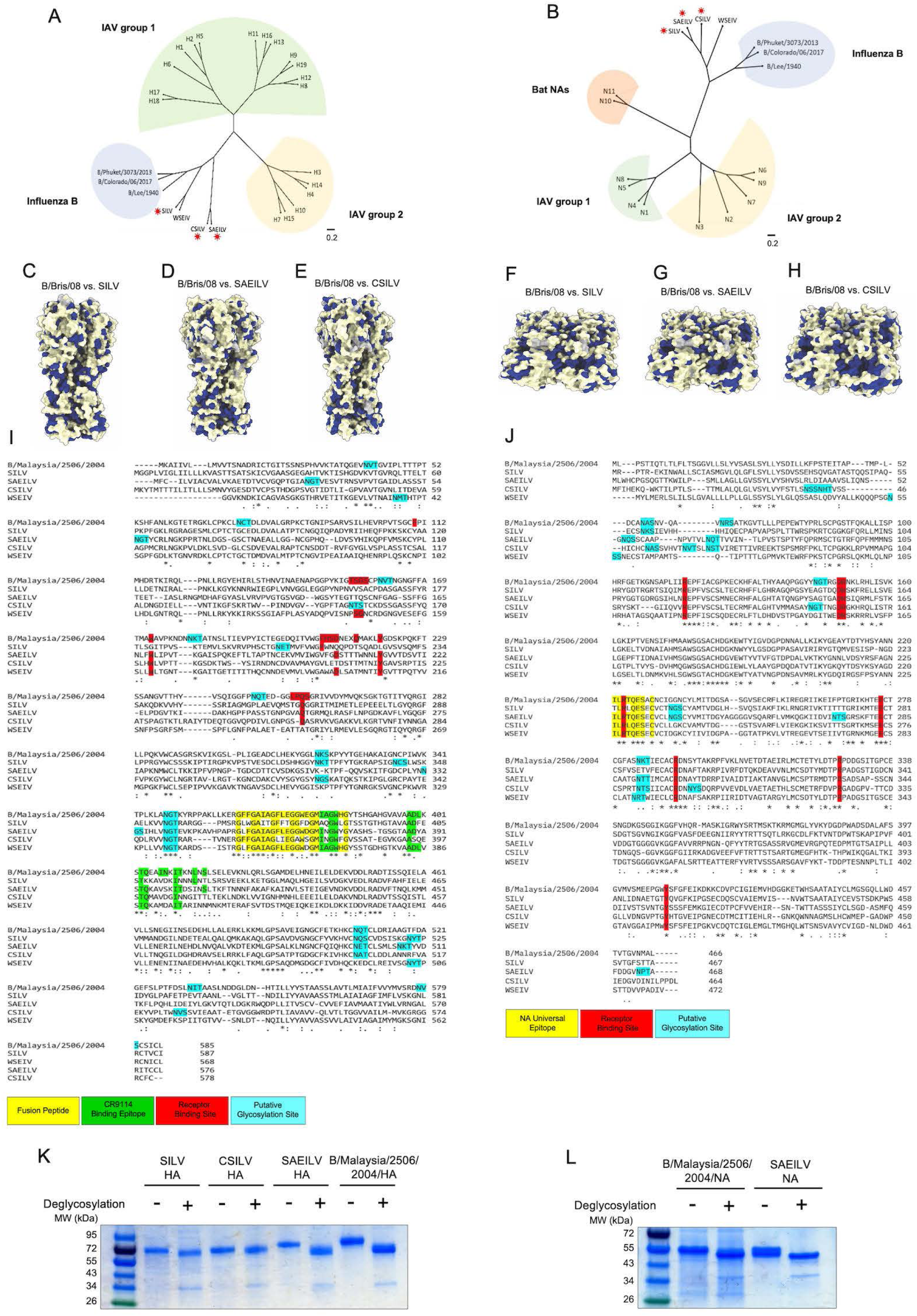
Phylogenetic and comparative sequence analysis of SILV, SAEILV and CSILV glycoproteins. Phylogenetic analysis of the salamander influenza-like virus (SILV), Siamese algae-eater influenza like virus (SAEILV) and chum salmon influenza-like virus (CSILV) HA and NA amino acid sequences and representative sequences from influenza A and B HAs (**A**) and NAs (**B**) were performed by the maximum likelihood method. The scale bar shows estimated amino acid substitutions per site. Conserved amino acid residues of the SILV HA and NA (**C**, **F**), SAEILV HA and NA (**D**, **G**), CSILV HA and NA (**E**, **H**) respectively relative to influenza B virus HA and NA are highlighted in blue on top of the publicly available structure of influenza B/Brisbane/60/2008 HA (PDB# 4FQM) and NA (PDB# 4CPL) by ChimeraX. Amino acid sequence alignment of the SILV HA and NA, SAEILV HA, NA and, CSILV HA and NA and WSEIV HA and NA are displayed against influenza B/Malaysia/2506/2004 HA and NA, (**I**) and (**J**). Key features are shown in different colors. Asterisks indicate identical amino acids. SDS-PAGE analysis of recombinant expressed SILV, SAEILV and CSILV HAs (**K**) and SAEILV NAs (**L**) and influenza B/Malaysia/2506/2004 HA and NA in deglycosylated and non-deglycosylated conditions.

It is worth noting that we were only able to produce SAEILV NA protein recombinantly in our baculovirus expression system. An N-terminal vasodilator stimulating phosphate (VASP) tetramerization domain was used to maintain the tetramerized structure of the head domain of NA. PNGaseF treatment of the B/Malaysia/2506/2004 NA and SAEILV NA did not reveal any overt differences in glycosylation patterns across the glycoproteins with comparable shifts in molecular weights pre- and post-deglycosylation (Fig.1L).

### Receptor specificity

To study the receptor binding specificity of these novel influenza B-like viruses, we performed a hemagglutination assay (HA assay) that exploits the ability of the HA to bind sialic acid receptors on red blood cells (RBCs), thereby cross-linking them and preventing gravity-induced pelleting(*29*). We expressed and tested the recombinant HA proteins (SILV, SAEILV and CSILV) as soluble trimers in the baculovirus system. Recombinant SILV failed to induce hemagglutination, while both SAEILV and CSILV caused hemagglutination to both chicken and turkey RBCs (Fig. 2A), suggesting that they can recognize canonical influenza virus receptors. The absence of hemagglutination was also noted for the recently identified influenza B-like WSEIV which binds to GM2 and for H17, H18 and H19 which use the major histocompatibility complex as receptor, raising questions about the receptor usage of the SILV HA(*14, 21, 22*). To investigate the receptor usage of these influenza B-like viruses more comprehensively, we used glycan microarrays containing synthetic glycans with α2,3- and α2,6-linked sialic acid (Neu5Ac) presented as linear structures, bi-and tri-antennary glycans including sialyl Lewis X (SLe^x^) structures(*17, 30*). The same approach had been previously employed to identify a sialylated ganglioside receptor for WSEIV(*14*). As expected, the B/Netherlands/2914/2015 control bound to both glycans containing α2,6-linked Neu5Ac or α2,3-linked Neu5Ac (Fig. 2E). However, we did not detect binding of the recombinant SILV HA to any of the glycans present on the microarray (Fig. 2B), which agrees with the lack of hemagglutination activity. We also did not detect any binding to glycolipids as we did for the WSEIV. SAEILV HA showed robust binding to α2,3-linked and to SLe^x^ structures and even appeared to bind some α2,6 Neu5Ac (Fig. 2C). CSILV HA had a more restricted binding repertoire and predominantly bound to α2,3-linked Neu5Ac (Fig. 2D). Interestingly CSILV apparently needs at least two α2,3-linked Neu5Ac within a *N*-glycan as only compounds **10** and **21** are bound and not compounds **6** and **17**, furthermore SAEILV was able to bind linear structures whereas CSILV was not. In contrast to SAEILV, CSILV was not able to accommodate SLe^x^, also not when presented as complex N-glycans as structures **11** and **22**. Conclusively all three HAs have a distinct receptor binding specificity with that of the SILV being an unknown, while SAEILV and CSILV both can bind canonical avian-type receptors

**Fig. 2.**
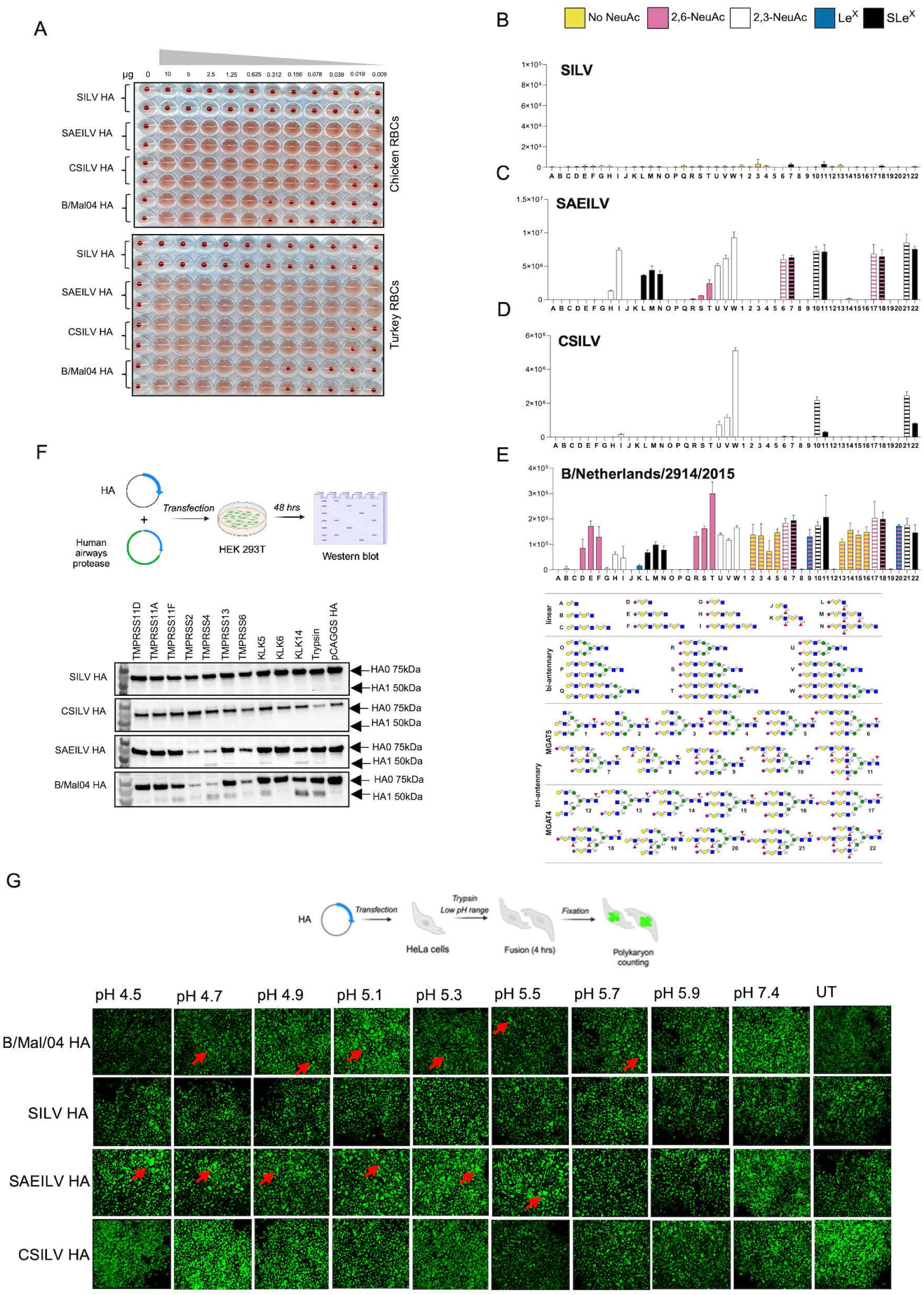
Functional profile of SILV, SAEILV and CSILV HAs. (A) Hemagglutination assays with recombinant HA proteins from salamander influenza-like virus (SILV), Siamese algae-eater influenza like virus (SAEILV) and chum salmon influenza-like virus (CSILV) were performed with turkey and chicken red blood cells (RBCs). Samples were added at an initial concentration of 10 μg/ml and serially diluted two-fold. (B-E). Sialic acid binding of recombinant HA proteins of SILV, CSILV and SAEILV was tested on a glycan microarray containing synthetic glycans with α2,3- and α2,6-linked sialic acid (Neu5Ac) presented as linear structures, bi-and tri-antennary glycans including sialyl lewis X (SLe^x^) structures. B/Netherlands/2914/2015 served as positive controls (**B-F**). Glycans A-M and O-W represent linear and bi-antennary structures respectively with indicated backbones. Glycans 1-22 represents tri-antennary *N*-glycans, either the MGAT4 or MGAT5 arm was elongated to two LacNAc repeating units. Glycans were terminating without a NeuAc (yellow), or with α2,6-linked NeuAc (pink), α2,3-linked NeuAc (white), Le^x^ (blue), or SLe^x^ (black). Bars with two colors indicate glycans terminating in different epitopes on different arms. Mean relative fluorescence unit (RFU) ± SD are shown. (**G**) Experiment setup: HEK 293T cells were co-transfected with two plasmids, one encoding the full length of corresponding HA and one encoding human airway proteases. The HA cleavage state was determined after a 48-hour transfection. Representative western blots show the bands of uncleaved HA0 and cleaved HA1. Cells transfected with pCAGGS HA, either untreated or exposed to TPCK-treated trypsin, served as assay controls. (**H**) Experiment setup: HeLa cells were transfected with pCAGGS plasmid expressing full length of corresponding HA and were treated trypsin, and exposed to an acidic buffer, and allowed to recover to determine for cell-to-cell fusion. (**I**) Representative photographs showing polykaryon formation in cells transfected with pCAGGS SILV, SAEILV, CSILV, and B/Malaysia/2506/2004 HA.

### Cleavage profile of HAs by human airways proteases

The activation of viral glycoproteins by host proteases is often necessary for viral fusion, infection, and pathogenicity. HA usually binds to sialyated glycans on the host cell surface from where the virus can then be transported into the cell via endocytosis. To gain membrane fusion competence, HA relies on host cell proteases for the cleavage of the HA0 precursor into the HA1 and HA2 polypeptides(*31*). A 2020 study investigated the differences in preferential proteolytic activation of influenza A and B viruses HAs by human respiratory epithelium enzymes(*32*). Influenza B virus HA0 has been found to be activated by a broader panel of type II transmembrane serine proteases (TTSP) and kallikrein (KLK) enzymes with much higher efficiency than influenza A virus HA0. Given the established involvement of TTSPs and KLKs in the cleavage and activation of influenza B viruses(*32*), we aimed to investigate the extent to which of these proteases were able to activate the HAs of the novel influenza B-like viruses. Human embryo kidney (HEK) 293T cells were co-transfected with pCDNA3.1 plasmids encoding human airway enzymes and pCAGGS expression plasmid encoding the corresponding full-length HA of SILV, SAEILV, CSILV, or B/Malaysia/2506/2004 HA (Fig. 2F). Cells transfected with only pCAGGS expression plasmid encoding the corresponding full-length HA served as an untreated control. As another control, pCAGGS HAs transfected cells were incubated briefly with N-tosyl-L-phenylalanine chloromethyl ketone (TPCK)-treated trypsin before harvesting the cells. 48 hours post transfection, the transfected cells treated, or mock treated, with TPCK-treated trypsin were harvested for detection of HA cleavage by Western blot. Cleavage was detected using polyclonal antisera generated in mice immunized with each corresponding recombinant HA. As evidenced by the presence of both an HA1 band (∼50 kDA) and an HA0 band (∼75 kDA), B/Malaysia/2506/2004 HA was cleaved by various human airways enzymes. No cleavage was observed in cells that were transfected with only the pCAGGS HA plasmid. Strikingly, none of the selected proteases or trypsin were able to cleave and activate the SILV and CSILV HA. On the other hand, SAEILV HA0 was cleaved by TMPRSS4, TMPRSS6, KLK5, KLK14 and trypsin.

### Activation pH of HAs from influenza B-like viruses

We next investigated whether HA0 cleavage translated into HA activation and subsequent membrane fusion using a polykaryon assay (Fig. 2G). HeLa cells transfected with pCAGGS plasmid expressing full length HA were treated with trypsin to activate the HA on the cell surface. Cells were then exposed to different pH conditions (4.5 to 5.9) for 15 minutes to promote membrane fusion. Cell-cell fusion was detected via the presence of polykaryons. Only cells transfected with SAEILV HA induced polykaryon formation at a variety of different pHs. Cells treated at a pH of 7.0 and untransfected cells did not form any syncytia, showing that the polykaryon formation was HA specific and pH dependent (Fig. 2I). Since SILV HA was not cleaved by trypsin and CSILV only to a low degree, it is not surprising that they did not show fusion activity in this assay. Their pH of fusion remains to be determined once appropriate proteases for their cleavage have been identified.

### The structural basis of influenza B-like HA receptor diversity

We were also interested in determining and comparing the structures of HAs of influenza B-like viruses with those of *bona fide* influenza B viruses. We first determined the cryo-electron microscopy (cryo-EM) structure of the previously described influenza B-like WSEIV HA at 2.67 Å resolution, and observed HA1 and HA2 subunits, glycans, RBS, and fusion peptide (Fig. 3A). Out of 3 predicted glycosylation sites in the WSEIV HA sequence, clear glycan densities were observed on residues N36 and N336 but not on the HA2 C-terminus-proximal residue N503 (Fig. 3A). The overall WSEIV HA trimer resembles the influenza B/Bris/08 virus HA (PDB 4FQM)(*25*), despite only sharing 44% amino acid sequence homology. There was a slight shift of angle in the central stem helix between WSEIV HA and B/Bris/08 HA, which led to a more drastic shift of the HA head (Fig. 3B). Surface charge analysis revealed distinct charge patterns on the HA head: WSEIV HA had negatively charged patches within and around the RBS, while B/Bris08 HA had positively charged patches within and around the RBS (Fig. 3C). In addition, the key residues within the RBS pocket of WSEIV HA are different from B/Bris/08 HA (Fig.3D). These discrepancies in the RBS also support different receptor preferences for these influenza B-like HAs.

**Fig. 3.**
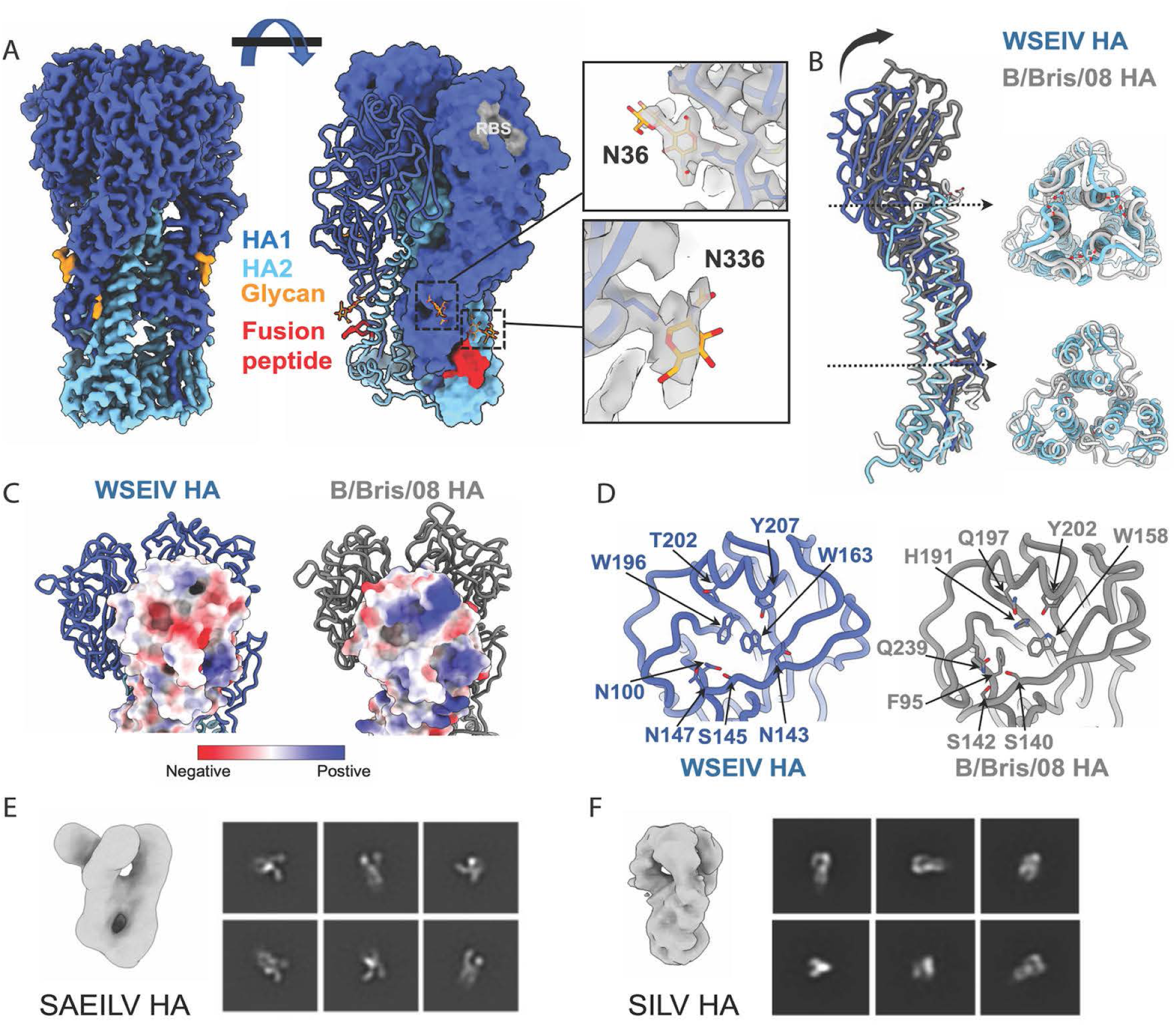
WSEIV HA cryoEM structure resembles influenza B HA. (**A**). CryoEM map and model of WSEIV HA showing the HA1/HA2 subunits, fusion peptide, RBS pocket, and N-linked glycans. (**B**). Models of WSEIV HA and B/Brisbane/60/2008 HA (PDB 4FQM) are aligned by the central helix of HA2. (**C**). The surface of WSEIV HA and B/Brisbane/60/2008 HA are colored by electrostatic potential using ChimeraX. (**D**). Key residues within the RBS of WSEIV HA and B/Brisbane/60/2008 HA. Ab-initio reconstruction maps and selected 2D classes of SAEILV HA (**E**) and SILV HA (**F**) from negative stain EM

Our attempts to obtain high resolution cryoEM structures of SAEILV/CSILV/SILV HA were unsuccessful. However, low resolution negative stain EM maps were acquired for SAEILV and SILV HA (Fig. 3E and F). The 3D-reconstruction maps show SILV HA resembles the canonical shape of influenza virus HA, while SAEILV HA has an open and flexible head domain. The unstable head domain in the recombinant HA protein makes structural determination difficult. Next, we predicted the models of the four influenza B-like virus HAs using the AlphaFold3 (AF3) server(*33*). The predicted models showed overall high confidence of accuracy on the protein backbones of the HA ectodomains, especially on the HA head regions, but the model quality of the cleavage sites and fusion peptides were quite poor (Fig. S1A). The AF3 predicted model of WSEIV HA highly resembles the cryoEM structure with a root mean square deviation (RMSD) of 1.298 Å (Fig. S1B). The predicted models also revealed divergent key residues located within the RBS pockets (Fig. S1C and Table. S1), which provide structural basis for their distinct receptor binding profiles (Fig. 2B-E).

### Neuraminidase activity and sensitivity to multiple neuraminidase inhibitors

NA is a fascinating protein that plays multiple essential roles in the influenza virus life cycle. Our efforts to express recombinant protein from SILV and CSILV NAs were unsuccessful, thereby hindering further functional and structural characterization of these newly identified influenza B-like virus NAs. Given that the primary active site residues that are crucial for the sialidase activity of the NA are conserved, we examined the functional activity of the SAEILV NA by performing an enzyme-linked lectin assay (ELLA) using fetuin as a substrate. Fetuin contains mono-, bi, and triantennary glycans with 2,3- and α2,6-linked sialic acids in a 2:1 ratio (*34*). Fetuin-coated 96 well plates were incubated with serially diluted proteins overnight at four different temperatures (4°C, 20°C, 33°C and 37°C) in order to determine the temperature dependent profile of the NAs. These temperatures were partially chosen to cover the range of temperatures that Siamese algae eater fish might experience in their freshwater habitat. NA enzymatic activity was quantified using peanut agglutinin (PNA) lectin which binds to exposed galactose residues that remain once terminal sialic acid is cleaved off. B/Malaysia/2506/2004 was used as a comparative control. We found that SAEILV NA has neuraminidase activity, and enzymatic activity is also slightly higher compared to B/Malaysia/2506/2004 NA at the tested temperatures (Fig. 4A). The inverse of the half-maximum lectin binding was used to calculate the specific activity of the NA in order to better understand the temperature dependency (Fig. 4B). As expected, we found that each NA’s activity gradually decreased at lower temperatures. Additionally, the pattern of specific activity profiles for the B/Malaysia/2506/2004 NA and the SAEILV NA were nearly the same.

**Fig. 4.**
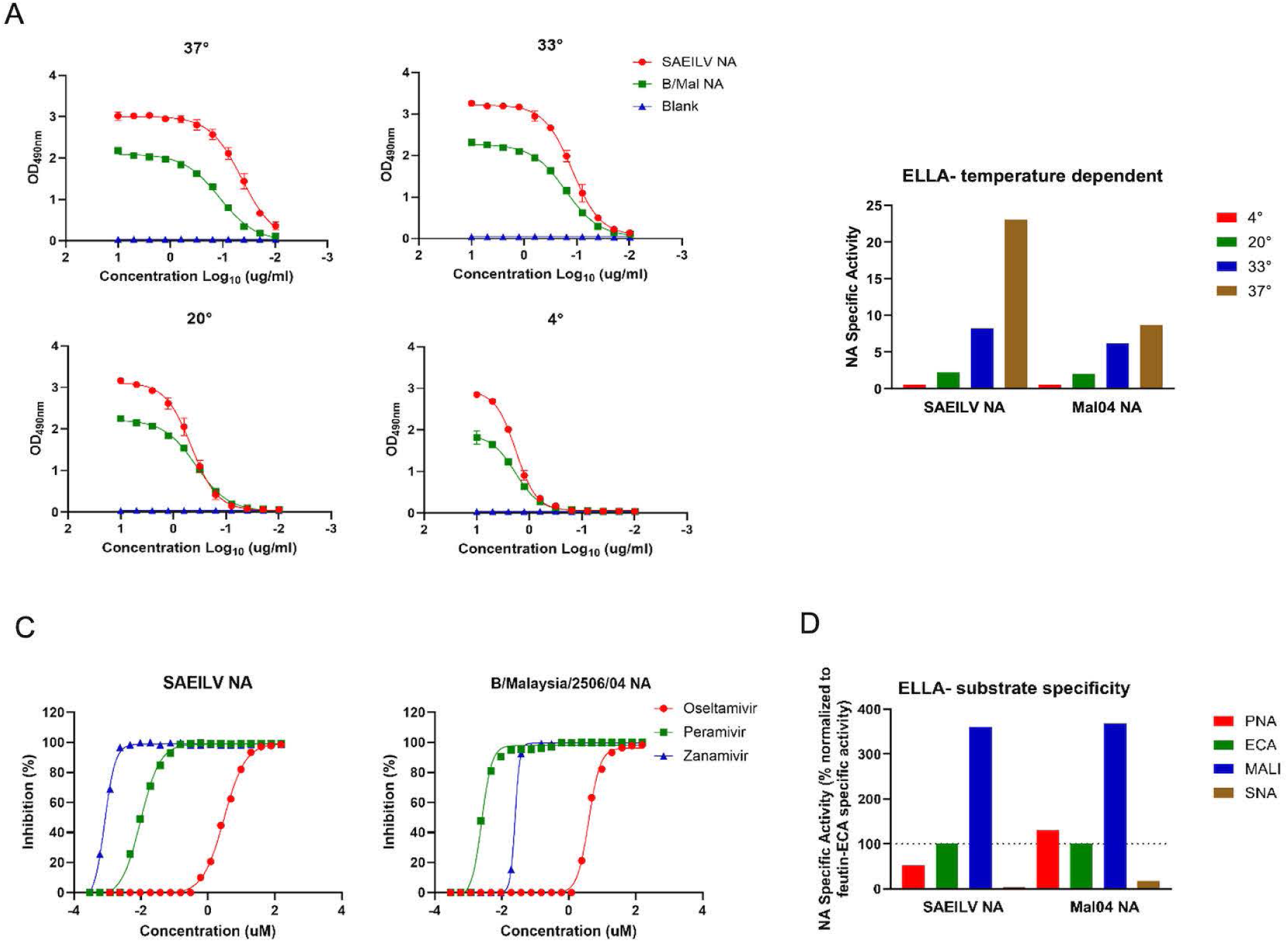
NA specific activities and cleavage specificities of the SAEILV NA. (**A**) Neuraminidase enzymatic activity of recombinant SAEILV and B/Malaysia/2506/2004 NA proteins was examined via enzyme linked lectin assay (ELLA) using fetuin, at four different temperatures: 4, 20, 33 and 37°C. The curves indicate absorbance measured at 490 nm with error bars indicating standard deviation. (**B**) Specific enzymatic activity (inverse of half-maximum lectin binding) is shown for each individual NA at each temperature. (**C**) Inhibitory susceptibility to three NA inhibitors (oseltamivir, peramivir and zanamivir) was evaluated in an ELLA-based neuraminidase inhibition assay. (**D**) Specific activity of SAEILV and B/Malaysia/2506/2004 NA as determined by ELLA using different glycoproteins-lectin combinations normalized to the fetuin-ECA.

To explore the susceptibility of the SAEILV NA to clinically approved NA inhibitors, the 50% inhibitory concentration (IC_50_) of oseltamivir, peramivir and zanamivir were determined using an ELLA-based neuraminidase inhibition assay. In concordance with previous findings for Wuhan spiny eel virus NA(*14, 23*), SAEILV NA was also more sensitive to peramivir and zanamivir than oseltamivir (Fig. 4C). These results also support our finding that residues within the enzymatic active site are highly conserved between influenza B virus NA and SAEILV NA, enabling NA inhibitor binding.

### Enzymatic activity of the SAEILV NA for differently linked sialic acids

Next, we examined the activities of the SAEILV NA for differently linked sialic acids using lectins with different binding specificities in a previously described ELLA protocol(*35*). Lectins with different binding specificities were used to determine if the NAs preferentially cleaved 2,3- and α2,6-linked sialic acid. *Erythrina crista-galli* lectin (ECA) and peanut agglutinin (PNA) bind to non-siaylated *N*- and *O*-linked sugars respectively and their binding would be in general higher in the presence of active NA (*36, 37*). *Maackia amurensis* lectin I (MAL I) and *Sambucus nigra* lectin (SNA) specifically bind α2,3- or α2,6-linked Neu5Ac, respectively, and their binding decreases with NA cleavage of the target substrates(*25, 38*). Specific activities (activity per amount of protein) of the SAEILV and B/Malaysia/2506/2004 NA proteins were determined for each glycoprotein-lectin combinations and plotted relative to the specific activity determined for fetuin-ECA. In line with the enzymatic activity, both proteins have equal substrate specificity and comparable cleavage profiles (Fig. 4D).

### The structural basis of NA active sites

To structurally compare the NA active site, we obtained a 2.75 Å cryoEM 3D reconstruction of the SAEILV NA head (Fig. 5A). We observed cryoEM density for one glycosylation site at position N281 located on the underside of NA (Fig. 5A). This glycan is also conserved in CSILV NA, previously reported WSEIV NA (PDB 7XVU)(*23*) as well as influenza B virus NA (Fig. 1J & Fig. 5B). Interestingly, despite the ∼ 40% sequence identity, the catalytic residues within the NA active site are highly conserved among SAEILV, WSEIV, and influenza B virus NA, with only one exception of a W407G mutation in SAEILV NA (Fig. 5B). The conserved residues in the NA active site pocket provide the structural basis of the susceptibility of SAEILV NA to neuraminidase inhibitors. On the contrary, the residues around the NA active site are less conserved as indicated by different surface charge patterns of SAEILV, WSEIV, and influenza B virus NAs (Fig. 5C), which also suggests their preference for substrates could be different. Additionally, we observed the potential density of a calcium ion in the cryoEM reconstruction map (Fig. 5D), indicating the activity of SAEILV NA might be Ca^2+^ dependent, like for seasonal influenza virus NAs. To understand why the broadly reactive NA antibody 1G01 does not bind SAEILV NA, we superimposed the structure of 1G01+ A/California/04/2009 (H1N1) NA (PDB 6Q23) (*38*) with SAEILV NA. There is an obvious clash between NA Y239 and the 1G01 HCDR3 loop. In addition, A/California/04/2009 N1 K150/K432 form additional salt bridges with 1G01, while SAEILV NA lacks such interactions (Fig. 5E). Like HA, we used AF3 to generate models from the sequences of CSILV and SILV NAs. Both models were predicted with >90% overall accuracy on the tetrameric NA head (Fig. S1D and E). The AF3 predicted SAEILV NA structure highly resembles our cryoEM structure with a RMSD of 0.752 Å on the NA protomer (Fig. S1F). Again, the key residues within the NA active site are highly conserved among these 3 NAs (Fig. S1G).

**Fig. 5.**
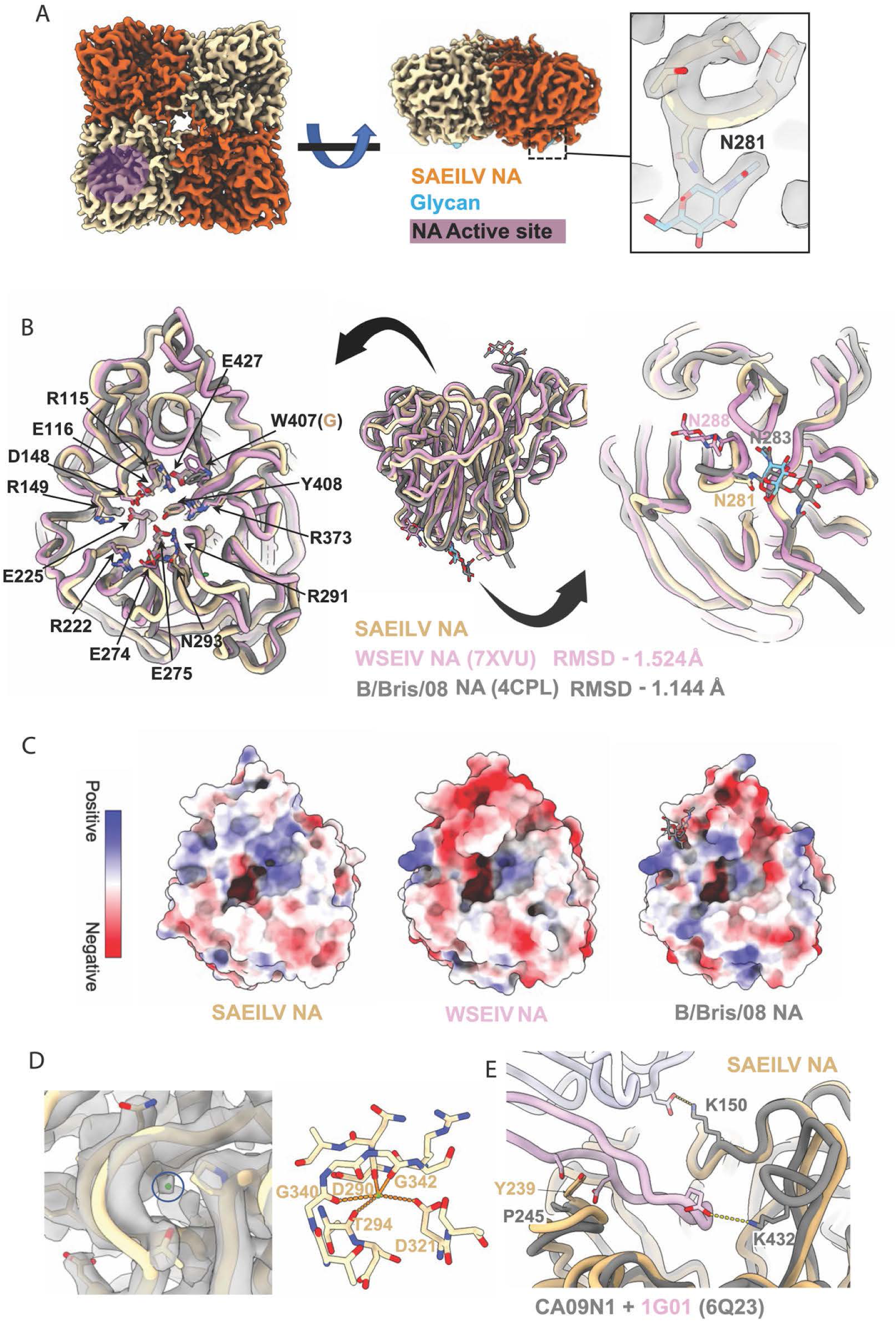
SAEILV NA cryoEM structure resembles influenza B NA. (**A**). CryoEM map of SAEILV NA showing the tetrameric NA head, active site, and N-linked glycan. (**B**). Aligned models of SAILV NA, WSEIV NA (PDB 7XVU), and B/Brisbane/60/2008 NA (PDB 4CPL) showing conserved secondary structures, key active site residues, and underside glycans among 3 NAs. (**C**). The surface and the active site pockets of SAEILV NA, WSEIV NA, and B/Brisbane/60/2008 NA are colored by electrostatic potential using ChimeraX. (**D**). CryoEM map and model showing the conserved calcium binding pocket in SAEILV NA. (**E**). Aligned models of SAEILV NA and 1G01 + A/California/04/2009 (H1N1) showing 1G01 HCDR3 clashes with SAEILV NA Y239.

### Characterizing the binding profile of mAbs using enzyme-linked immunosorbent assay (ELISA) and immunofluorescence

Next, we focused on characterizing the binding of broadly protective monoclonal antibodies to these novel influenza B-like glycoproteins (Fig. 6A, B, C). We used a panel of broadly cross-reactive human and mouse monoclonal antibodies which had been earlier characterized to bind to diverging lineages of influenza B virus(*25, 38–40*). To do so, ELISA and flow cytometry based cellular assays were performed and robust binding for most of the tested human and mouse monoclonal antibodies was observed for B/Malaysia/2506/2004 HA and NA which served as our experimental control. We did not detect any binding to SILV, SAEILV and CSILV HAs by any of human and mouse monoclonal antibodies. Only a positive control antibody targeting the his-tag on the recombinant proteins produced a signal, showing that the proteins were actually coated on the plates. As aforementioned, our efforts to express recombinant protein from SILV and CSILV NAs were unsuccessful, and we took two different approaches to characterize the binding profiles of these NAs with panel of human and mouse mAbs. Out of the panel of human and mouse mAbs used to probe SAEILV NA (which we could produce as recombinant protein), only one antibody, 2E01, showed low but detectable binding in ELISA, flow cytometry-based assay and in an immunofluorescence assay (IF assay) (Fig. 6B, C, S1C). Our group earlier showed anti-influenza B NA-mAb 2E01 demonstrated remarkable NA inhibition and binding activities against viruses belonging to the B/Yamagata/16/88-like and B/Victoria/2/87-like lineages and the ancestral B/Lee/1940 strain, which cumulatively span more than 70 years of antigenic drift.(*41*) A positive control in immunofluorescence assay, mouse antisera generated against SAEILV NA by vaccinating mice with recombinant SAEILV NA protein, was used. This serum indeed detected SAEILV NA expression on transfected cells. We did not detect any binding in flow cytometry or IF for SILV or CSILV NAs, but since we did not have specific sera as positive controls, we can also not prove that the transfected cells actually expressed these NAs. 4F11 (anti-influenza B virus NA) and 4C2 (anti-influenza B virus HA) mouse mAbs were used as negative control for the assay against the respective HAs and NAs and showed no signal as expected.

**Fig. 6.**
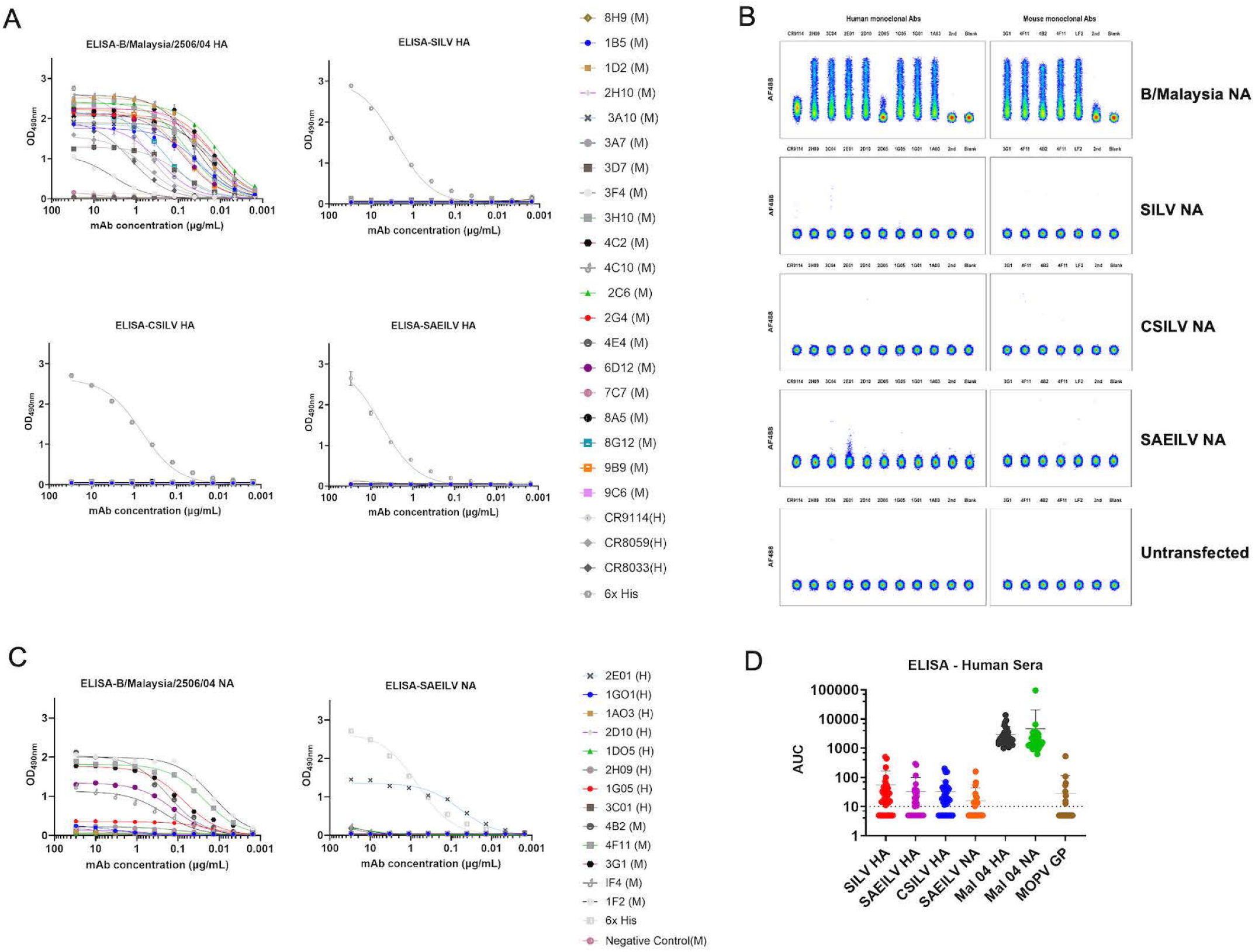
Binding profiles of the anti-influenza B virus HA and NA monoclonal antibodies (mAbs). (**A** and **C**) Broadly cross-reactive anti-influenza B virus HA and NA human and mouse monoclonal antibody binding to recombinant protein (SILV, SAEILV, CSILV and B/Malaysia/2506/2004 HA) and (SAEILV and B/Malaysia/2506/2004 NA) in enzyme linked immunosorbent assay. (**B**) Anti-influenza B virus human and mouse monoclonal antibody binding to cell surface expressed NAs of SILV, SAEILV, CSILV and B/Malaysia/2506/2004 by flow cytometry. (**D**) Cross reactive antibody profile of sera from post-seasonal influenza vaccination recipients. Antibodies against SILV, SAEILV and CSILV HA and SAEILV NA were determined via ELISA. Mopeia virus glycoprotein was used as negative control for baseline establishment.

Finally, to determine if humans have immunity to these HAs and NAs, we performed ELISAs with a panel of human sera (Fig. 6D). A glycoprotein from an arenavirus, Mopeia virus, to which humans are naïve, was used as negative control for baseline establishment. No reactivity above baseline was found against the HAs and NA of influenza B-like viruses (while reactivity to *bona fide* HA and NA of influenza B virus was high as expected), further supporting the finding that there is limited conservation of cross-reactive epitopes as also shown by our panel of mAbs.

## Discussion

Influenza viruses are well known to infect a broad spectrum of host species, however, there have been few studies of influenza-like viruses from fish. In a recent study, NA and HA sequences were identified in gills and respiratory tissue samples from salamander as well as Siamese algae and chum salmon fishes(*15*). Here, we report the functional and structural characterization of HA and NA of influenza B-like viruses from fish. The identification of novel influenza viruses in fish that are genotypically and phenotypically distinct from other influenza A viruses and clusters close to influenza B viruses suggest the occurrence of prolonged virus-host co-divergence with several host-switching events over time(*13, 15*). We found that, like the previously described WSEIV HA(*14*), SILV HA does not bind to canonical human or avian receptors. This lack of canonical receptor binding is likely due to specific structural features of the putative receptor binding site of SILV and WSEIV HA. The WSEIV HA structure contains a smaller RBS pocket with negatively charge patches within and around the RBS; this may result in steric or charge constraints that limit canonical α2,3- and α2 -linked Neu5Ac binding. CSILV and SAEILV on the other hand do interact with canonical avian-type receptors and are different from each other in their ability to interact with SLe^x^

Notably, SILV, SAEILV, CSILV and WSEIV NA show the highest similarity to influenza B virus NA with amino acid sequence identity of 44%, 40%, 34% and 48% respectively. While this is a relatively low sequence identity, these NAs display similar overall structures to other typical influenza virus NAs with a conserved active site. Our data clearly showed that SAEILV NA has similar neuraminidase activity as typical influenza B virus NAs and is sensitive to multiple NAIs like zanamivir, oseltamivir and peramivir, which is in concordance with the previous studies on WSEIV NA(*14, 23*). In addition, we found limited antigenic conservation between these influenza B-like viruses and *bona fide* influenza B viruses. No antibody-based immunity was detected against the HA and NA of these influenza B-like viruses in humans.

In this study, we resolved the high-resolution structures of WSEIV HA and SAEILV NA by cryoEM and performed nsEM 3D reconstructions of SAEILV HA and SILV HA. However, we were unable to observe the trimeric CSILV HA by nsEM, suggesting the recombinant CSILV HA is unstable and might exist in a monomeric form. AF3 predicted models were highly consistent with our WSEIV HA and SAEILV NA cryoEM structures, suggesting that the AF3 models of the remaining HAs are reliable preliminary models that can be used to facilitate future attempts to further stabilize these proteins for recombinant expression.

Seasonal epidemics caused by influenza B viruses were responsible for up to 52% of influenza-associated pediatric mortality during the last fifteen years(*42, 43*). So, influenza B virus surveillance and diagnostics in non-human hosts are critically important and improve our understanding of influenza B virus ecology and diversity. Looking at influenza B-like viruses from other animal lineages and vertebrate classes will add to our understanding of the origins and evolution of this important group of viruses.

In summary, the data from our structural and functional characterization of the novel influenza B-like viruses from fish reveal that the SILV HA is functionally divergent from influenza B virus HA. The SILV HA does not induce hemagglutination with chicken or turkey RBCs, nor does it bind to canonical human or avian receptors. Conversely, CSILV and SAEILV on the other hand do interact with canonical avian-type receptors. Additionally, SAEILV was activated by human airway enzymes and is fusogenic at a wide range of pH conditions. No cross-reactivity antibodies were found in the human serum suggesting that humans are immunologically naïve to these glycoproteins. We demonstrated that SAEILV NA protein has canonical overall structure and functions compared to other typical influenza virus NAs and is sensitive to all currently used NAIs. Our detailed structural studies of WSEIV HA and SAEILV NA provide insights into understanding the newly identified influenza B-like virus HA and NA and understanding the mechanism of lack of binding to pan NA mAb 1G01.

## Materials and methods

### Cell and recombinant proteins

Sf9 cells (CRL-1711, ATCC) used for baculovirus rescue were cultured in *Trichoplusia ni* medium-formulation Hink insect cell medium (TNM-FH, Gemini Bioproducts), supplemented with 10% fetal bovine serum (FBS, Gibco) and a penicillin (100 U/mL)-streptomycin (100 μg/mL) solution (Gibco). High Five cells (BTI-TN-5B1-4 subclone; Vienna Institute of Biotechnology)(*44*), utilized for protein expression, were grown in serum-free Sf-900 medium (Gibco) supplemented with penicillin (100 U/mL)-streptomycin (100 μg/mL) solution (Gibco). Human embryonic kidney (HEK) 293T (CRL-3216, ATCC) cells were propagated using Dulbecco’s modified Eagle’s medium (DMEM; Gibco) supplemented with 10% FBS, penicillin (100 U/mL)-streptomycin (100 μg/mL) solution and 1% 1 M 4-(2-hydroxyethyl)-1-piperazineethanesulfonic acid (HEPES). HeLa cells (CCL-2, ATCC) were grown in Eagle’s minimum essential medium (EMEM) supplemented with 10% FBS and penicillin (100 U/mL)-streptomycin (100 μg/mL) solution.

Recombinant HA and NA proteins were expressed in the baculovirus expression system, as described in previous reports(*26*). Ectodomains from SILV HA (Met 1-Gly 529), SAEILV HA (Met 1-Leu 515), CSILV HA (Met 1-Val 521) and WSEIV HA (Met 1-Thr 522) were cloned into modified pFastBacDual transfer plasmids containing a T4 trimerization domain with a thrombin cleavage site and a hexhistidine purification tag (His-tag). Coding sequences from SILV NA (Ser 79- Ala 467), SAEILV NA (Ser 82- Ala 468), CSILV NA (Ser 81- Leu 464) were cloned also into modified pFastBacDual transfer plasmids which included a signal peptide, a hexhistidine purification tag (His tag), a vasodilator stimulating phosphoprotein (VASP) tetramerization domain and a thrombin cleavage site. The recombinant pFastBac constructs were introduced into DH10Bac bacteria (Invitrogen). Transformed colonies were selected, screened, and cultured for the extraction of bacmid DNA. This bacmid DNA was used to transfect the Sf9 cells to rescue recombinant baculovirus and then used to infect High Five cells for protein expression. Supernatants were subsequently collected by low-speed centrifugation approximately 72 hours post infection and protein was purified by Ni^2+^-nitrilotriacetic acid resin chromatography (Qiagen).

### Phylogenetic and sequences analyses

Representative HA and NA amino acid sequences of influenza A and B viruses for the phylogenetic trees were obtained from the Global Initiative on Sharing All Influenza Data (GISAID) and GenBank (H1-H18: EPI1349891, EPI899625, EPI673678, EPI1007628, EPI942074, EPI1154383, EPI1090164, EPI1154159, EPI1103524, EPI953583, EPI774886, EPI1007631, EPI967018, EPI750076, EPI965435, EPI939704, EPI356309, EPI486922; B/Lee/1940 HA: EPI243230; B/Phuket/3073/2013 HA: EPI1799824; B/Colorado/06/2017 HA: EPI969380; SILV HA: QOE76814.1; SAEILV HA: QOE76806.1; CSILV HA: QOE76830.1; WSEIV HA: AVM87624.1. N1-N11: EPI1381203, EPI899627, EPI939823, EPI1154448, EPI1007658, EPI939830, EPI750078, EPI941550, EPI965439, EPI356311, EPI356298; B/Lee/1940 NA: EPI366432; B/Phuket/3073/2013 NA: EPI1799823; B/Colorado/06/2017 NA: EPI969379; SILV NA: QOE76813.1; SAEILV NA: QOE76806.1; CSILV NA: QEO76829.1; WSEIV NA: AVM87625.1). Amino acid sequences were aligned using multiple sequence comparison by log-expectation (MUSCLE), and a phylogenetic tree was constructed by the maximum likelihood method with the MEGA11 software(*45*) employing default parameters and 1,000 bootstrap replications. The tree was cleaned and edited using FigTree v1.4 (http://tree.bio.ed.ac.uk/software/figtree/) and labels along with highlights were added in Microsoft PowerPoint. Pairwise alignment of the SILV HA and NA, SAEILV HA and NA CSILV HA and NA, WSEIV HA and NA and B/Malaysia/2506/2004 HA (ACR15732.1), B/Malaysia/2506/2004 NA (ACR15736.1) was carried out using Clustal Omega V1.2.4. Potential N-glycosylation sites (PGS) of the HA and NA were predicted using the NetNGlyc 1.0 Server (https://services.healthtech.dtu.dk/services/NetNGlyc-1.0) on the basis of the sequon motif N-X-S/T, where X can be any amino acid except proline. Identical residues were indicated by asterisks. Functionally and antigenically relevant features have been annotated.

### Deglycosidase treatment and SDS-PAGE

In step 1, 5 μg of recombinant protein was mixed with 1μL of deglycosylation mix buffer 2 (NEB), and sufficient PBS was added to achieve a total volume of 8 μL. The mixture was heated at 75°C for 10 minutes to denature the glycoproteins and then chilled on ice. In step 2, 1μL protein deglycosylation mix II was added to the non-control tube and mixed gently, while 1μL PBS was added to control tube. The reaction mixtures were incubated at room temperature (RT) for 30 minutes, followed by incubation at 37°C for 16 hours. Subsequently, the samples were treated with 2X Laemmli loading buffer (BioRad), supplemented with 10% beta-mercaptoethanol. The samples were then heated for 10 minutes at 98 °C prior to loading them onto sodium dodecyl-sulfate polyacrylamide gel (4–20% Mini-PROTEAN^®^TGX™ Precast Protein Gels, BioRad) afterwards were subsequently stained with Coomassie blue fast stain solution for 1 hour, shaking at RT. The staining solution was removed, and the gel was de-stained following incubations with distilled water.

### Hemagglutination assay

Ten μg of recombinant HA were serially diluted 2-fold in phosphate-buffered saline (PBS) and incubated with 0.5% chicken or turkey red blood cells (RBCs) in a total volume of 100 μL/well for 45 min at 4°C. Hemagglutination of RBCs was visually assessed.

### Glycan array

Glycan array binding analysis of the rHAs was conducted as previously described (*17, 30, 46*). Briefly, recombinant hexahistidine-tagged HA was pre-complexed with a mouse anti-his Alexa 647 (Thermo Fisher Scientific, catalog no. MA1-21315-A647) and goat anti-mouse Alexa 647 antibodies (Invitrogen, catalog no. A21235). This preparation was performed in 50 μL PBS-T (phosphate-buffered saline with 0.1% Tween-20) at a 4:2:1 molar ratio, incubated for 15 minutes on ice, and the applied on the array for 90 minutes. After multiple washes with PBS-T, PBS, and deionized water, the arrays were scanned to detect HA binding using an Innoscan 710 (Innopsys). The B/Netherlands/2914/2015 strain was a kind gift of Dr. Ron Fouchier (Erasmus Medical Center, Rotterdam, The Netherlands) and was detected using CR9114 and goat anti-human Alexa 647 antibodies (Invitrogen, catalog A-21445). Mean relative fluorescence unit (RFU) and standard deviation values were imported into Prism 7.0 and the corresponding graph was generated.

### Protease cleavage assay

To assess HA0 cleavage in protease-expressing cells, HEK293T cells were seeded onto a poly-D-lysine-coated twelve-well plate at a density of 300,000 cells per well. The following day, cells were co-transfected with 0.5 ug of the pCAGGS expression plasmid encoding the corresponding full-length HA and 0.5 ug of pCDNA3.1 plasmids encoding human airway proteases (Genescript) using the TransIT-LT1 transfection system (Mirus Bio). At 48 hours post transfection, the control well was exposed to 5 ug/ml of TPCK-treated trypsin for 15 min at 33°C. Cells were lysed with radio-immunoprecipitation assay (RIPA) buffer (ThermoFisher, USA) in presence of phosphatase/protease inhibitors cocktail (ThermoFisher, USA). The lysate was centrifuged (15 minutes at 4 °C and 17,005 × *g*) and the supernatant was collected. Western blot performed with standard protocol. Briefly, proteins were separated on a 4–20% Mini-PROTEAN^®^TGX™ Precast Protein Gels, (BioRad) and transferred on a nitrocellulose membrane using the iBlot transfer and stack device (Invitrogen) at 20 V for 7 min. The membranes were blocked in 5% nonfat dry milk (Bio-Rad) in Phosphate buffered saline with 0.5% Tween-20 (PBS-T) for 1 h at RT with shaking. The blocking solution was removed and corresponding anti-HA antisera (generated in-house by immunizing female BALB/c mice intramuscularly with SILV HA, SAEILV HA and CSILV HA recombinant protein) diluted 1:2000 in PBS supplemented with 1% nonfat dry milk was added. The membranes were incubated overnight and washed three times with PBS-T. The following day, anti-mouse IgG-horseradish peroxidase linked (1:5000; cat. No 7076; cell signaling technology, Inc) was added for 1 h at RT and the membranes were washed. The membranes were developed by adding ECL prime and incubated for 5 min. Imaging was performed with a ChemiDoc^™^ MP Imaging System (Bio-Rad).

### Cell–cell fusion assay

To investigate the impact of stabilization of the pH switch regions on HA fusogenicity, we performed cell–cell fusion experiments as previously described (*32*). HeLa cells were seeded in 96-well tissue culture plate at density of 20,000 cells per well. The next day cells were transfected with a pCAGGS plasmid expressing the full-length HA using the TransIT-LT1 transfection system (Mirus Bio). After 24 hours, the growth medium was replaced with 0.2% FBS, and cells were further incubated at 33°C. The following day, cells were washed with Dulbecco’s phosphate buffered saline (DPBS) and treated with 5 μg/mL TPCK-treated trypsin for 10 minutes at 33°C before being exposed to pH-adjusted DPBS, ranging from pH 4.5 to 5.9, adjusted with citric acid. After a 15 min acid pulse, the cells were washed twice with DPBS and subsequently incubated with DMEM supplemented with 10% FBS and penicillin-streptomycin (Pen-Strep) for 4 hours at 33°. The cells were fixed with 3.7% (v/v) paraformaldehyde (PFA) and permeabilized with 0.1% Triton X-100 for 15 minutes each. Staining was performed using the HCS cellMask Green (Life Technologies) as per manufacturer’s instructions for 30 min at 5 μg/mL. Images were acquired usingEVOS M5000 (ThermoFisher Scientific) fluorescence microscope.

### ELISAs

Ninety-six well flat bottom Immulon 4 HBX plates (Catalog no. 3855; Thermo Scientific) were coated for overnight at 4°C with 2 μg/mL of purified recombinant protein at 50 μL/well in PBS. The next day, the plates were washed using PBS-T, 225 μL/well, using a Biotek 405 Microplate washer. Plates were blocked using 3% milk protein in PBS-T for 1 hour at RT. The antibodies were diluted to a starting concentration of 30 μg/mL, serially diluted 1:3 in blocking solution, and incubated for 2 hours at RT. For human sera ELISAs the sera were diluted with initial dilution 1:100 with two-fold serial dilutions and incubated at RT for 2 h. The microtiter plates were washed three times with PBS-T and 50 μL of goat anti-human IgG (Fab specific) horseradish peroxidase antibody (HRP; Sigma, #A0293) or goat anti-mouse horseradish peroxidase antibody (Rockland; 610-1302) diluted 1:3000 in blocking solution was added to all wells and incubated for 1 h at RT. The plates were washed three times and 100 μL SigmaFast o-phenylenediamine dihydrochloride (OPD; Sigma) was added to all wells. After 10 min, the reaction was stopped with 50 μL 3 M hydrochloric acid (Thermo Fisher) and the plates were read at a wavelength of 490 nm with a plate reader (BioTek). The data were analyzed in Microsoft Excel and GraphPad Prism 7. The data were visualized as binding curves by applying a nonlinear fit.

### Immunofluorescence microscopy

HEK 293T cells were plated at 3 × 10^5^ cells/mL in a sterile, 96 well plate and incubated overnight at 37°C with 5% CO_2_ as described previously (*47*). The following day, cells were checked for >99% confluence and washed with 1X PBS. Cells were transfected with a pCAGGS plasmid expressing the full-length NA using the TransIT-LT1 transfection system (Mirus Bio). After 24 hour the growth medium was replaced with DMEM supplemented 0.2% FBS, and cells were further incubated at 33°C for another day. The following day, cells were fixed with 3.7% PFA in PBS for 1h at RT. Next, the PFA was removed, and the cells were then washed two times with PBS and blocked with 3% milk in PBS for 1 hour at RT. The blocking solution was then removed and primary mAbs were added to their respective wells at a concentration of 10 ug/mL in 1% milk in PBS. Primary antibodies were incubated with shaking for 1 hour at RT. Plates were then washed 3 times with 1X PBS and goat anti-mouse Alexa Flour 488 conjugated antibody (Invitrogen) or goat anti-human Alexa Flour 488 (Invitrogen) and 4’, 6-diamidino-2-phenylindole (DAPI) both at 1:500 in 1% milk added to the cells for secondary staining for 1 hour. The plates were washed 3 times with PBS and then imaged using the EVOS M5000 (ThermoFisher Scientific) fluorescence microscope. Overlays between red and blue channels were made in ImageJ.

### Flow Cytometry for cell surface staining

In 96 well HEK 293T cells were transfected with a pCAGGS plasmid expressing the full-length NAs as explained above in immunofluorescence assay. Cells were collected after 48h transfection and were washed once with 2% FBS (HyClone) in phosphate buffered saline (PBS) (Corning, DPBS 1X (without calcium and magnesium)). Next, cells were incubated with 10 ug/mL of primary mAbs in V-bottom 96-well plates (Greiner) at 4 °C for 30 min. After washing with 2% FBS in PBS cells were incubated with and goat anti-mouse Alexa 488 conjugated antibody (Invitrogen) or goat anti-human Alexa 488 (Invitrogen) secondary antibody at 4 °C for 30 min in the dark. The samples were washed and re-suspended in PBS. Flow cytometric data were acquired on a FACScanto II flow cytometer. At least 50,000 events for each sample were recorded and data were analyzed with the FlowJo 10 software.

### Enzyme-linked lectin assay (ELLAs)

ELLAs, used to determine enzymatic activity of the NAs and their sensitivity to inhibition, were performed as described in detail in previous reports(*14, 35*). Flat-bottom Immulon 4HBX microtiter plates (Catalog no. 3855; Thermo Scientific) were coated with 25 ug/mL of fetuin (Sigma) diluted in 1x PBS, at 100ul per well and incubated overnight at 4°C. The next day, the fetuin coated plates were incubated with 2-fold dilutions of recombinant NAs starting at 10 ug/mL in sample diluent buffer (1x DPBS supplemented with 1% BSA, and 0.5% Tween 20) overnight. In order to determine the temperature-dependent profile of the NAs, the plates were incubated overnight at either 4, 20, 33, or 37°C. The following day, the plates were washed six times with PBS-T. Peanut agglutinin conjugated with HRP (PNA-HRP; Sigma) was diluted to 5 ug/mL in a conjugate diluent buffer (1x DPBS supplemented with 0.1% BSA) and added to washed plates. The plates were incubated for 2 h in the dark at RT. The plates were washed three times with PBST and developed with 100 μL of SigmaFast o-phenylenediamine dihydrochloride OPD.

To evaluate the potency of NA inhibitors on SAEILV NA, an NA inhibition assays using oseltamivir, peramivir or zanamivir were performed. Plates were coated overnight with fetuin as described above. In a separate U-bottom plate, inhibitor solutions with a starting concentration of 156.28 μM were serially diluted 2-fold in 1x PBS and were pre-incubated with the 15 μg of recombinant NAs for 1 hour on a shaker at RT. The fetuin coated plates were washed three times with PBS-T as described above. The inhibitor-protein dilutions were transferred to the fetuin plates and incubated at 37°C for 18 hours (overnight). The following day, the ELLA procedure was performed as described above. Substrate specificity characterization and specific enzyme activity determination was performed using a previously described ELLA assays [24]. In brief, 25 ug/mL of fetuin coated plates were incubated with serial dilutions of recombinant NA proteins sample diluent buffer. After the overnight incubation at 37°C, the plates were washed and incubated with either biotinylated ECA (1.25 μg/ml; Vector Laboratories), biotinylated PNA (2.5 μg/ml; Vector Laboratories), biotinylated SNA (1.25 μg/ml; Vector Laboratories), or biotinylated MAL I (2.5 μg/ml; Vector Laboratories). The binding of ECA, PNA, SNA, and MAL I was detected using horseradish peroxidase (HRP)-conjugated streptavidin (Thermo Fisher Scientific). Absorbance measurements were carried out at 490 nm following the development with SigmaFast OPD and specific enzyme activity (inverse of half-maximum lectin binding) was determined following a nonlinear regression analysis in GraphPad Prism 7 and was plotted normalized to the fetuin -ECA to determine activity per amount of protein in context of cleavage of α2,3 or α2,6-linked sialic acids.

### Human serum samples

The serum samples were obtained from an observational study approved by the Mount Sinai Hospital Institutional Review Board, IRB-16-00772. All study participants provided written consent before the biospecimens, and data were collected. All specimens were coded before processing and analysis. sera were stored at -80°C until analysis.

### Negative stain electron microscopy (nsEM)

Purified recombinant HA or NA were deposited on glow-discharged (PELCO easiGlow, Ted Pella, Inc.) carbon-coated 400 mesh copper grids (Electron Microscopy Sciences) at a concentration of approximately 20 μg/mL. Excess sample was blotted with filter paper. The grids were stained two times with 2% w/v uranyl formate for 10 s and 60 s respectively. Excess stains were blotted with filter paper. Grids were imaged on either a 200 kV Talos (Thermo Scientific) or a 120 kV Tecnai Spirit T12 (FEI) with an Eagle charge-coupled device (CCD) 4k camera (FEI). Images were collected at 72,000 or 52,000× magnification with pixel sizes of 2 Å and 2.06 Å, respectively. Micrographs were acquired using the EPU (Thermo Scientific) or Leginon(*48*) and data was processed with Appion(*49*) software package or CryoSPARC 4.0.(*50*)

### CryoEM grid preparation and imaging

Three uL purified recombinant HA or NA (0.4-0.8 mg/mL) were mixed with 0.5 mL 0.7% (w/v) Octyl-beta-Glucoside (OBG) before applied to glow-discharged grids (Electron Microscopy Services Cu 1.2/1.3 300-mesh). The grids were vitrified with Mark IV Vitrobot (Thermo Scientific) with following settings: 4°C, 100% humidity, 0 s wait time, 4.5-5.5 s blot time, blot force 1. After freezing, cryo grids were loaded into the 300 kV FEI Titan Krios (for WSEIV HA) or the 200 kV Glacios (Thermo Scientific) (for SAEILV NA), which were equipped with K3 Summit direct electron detector cameras (Gatan) and TFS Falcon IV (Thermo Scientific) respectively. Data were collected with approximate exposures of 50 e^−^/Å^2^ at magnifications of 105,000X or 190,000X for the Krios or Glacios, respectively. Data collection was automated using Leginon (*48*)or EPU. Further details are described in Table. S2.

### CryoEM data processing

Image pre-processing was performed with the Appion software(*49*) package or CryoSPARC Live. Micrograph movie frames collected from Krios were aligned, dose-weighted using the UCSF MotionCor2 software, and GCTF(*51*) was estimated. All micrographs were processed by CryoSPARC v4.0 for particle picking and reference-free 2D classification. Initial 2D classes of high quality were used as templates for template picking of datasets followed by 2D classifications to remove bad particles. The final maps of HA and NA were generated by non-uniform refinement with C3/C4 symmetry, followed by Global CTF refinement and another round of non-uniform refinement. More details are described in Table. S2.

### Atomic model building and refinement

AF3 predicted structures were used as the initial WSEIV HA and SAEILV NA models for refinement. Both models were manually fit into density using ChimeraX(*52*) followed by iterative manual model building in Coot(*53*) followed by refinements with Phenix(*54*) and Rosetta.(*55*) All the figures of maps and models were generated using ChimeraX.

### Data Analysis

Statistical analyses were performed using Prism 9 software (GraphPad). IC_50_ and area under curve (AUC) values were calculated using nonlinear regression (4 parameters) based on log_10_ transformed protein concentrations.

## Acknowledgements

We would also like to thank Edward Holmes at The University of Sydney for inspiring us to perform this study and for making the sequence information available. We appreciate the study participants of PARIS cohort, for generosity and willingness to enroll and donate samples for our research purpose.

## Author contributions

Conceptualization: GS, FK

Methodology: GS, JHu, DB, KV, R.P.d.V, GJB, VS and Jha

Investigation: GS, JHu, DB, KV, R.P.d.V, and Jha

Visualization: GS, FK, JHu and JHa

Supervision: JHa, ABW and FK

Writing—original draft: GS, JHu and FK.

Writing—review & editing: All authors contributed to the final manuscript version

## Data availability

All data are available in the manuscript or the supplementary materials. EM maps and models have been deposited in the EMDB and wwPDB, see Table S2. Reagents and antigens described in the manuscript can be provided by the Krammer laboratory pending scientific review and a completed material transfer agreement (and any required shipping/handling permits for viruses). Requests for the reagents and antigens should be submitted to.

## Supplementary Materials for

**Fig. S1.**
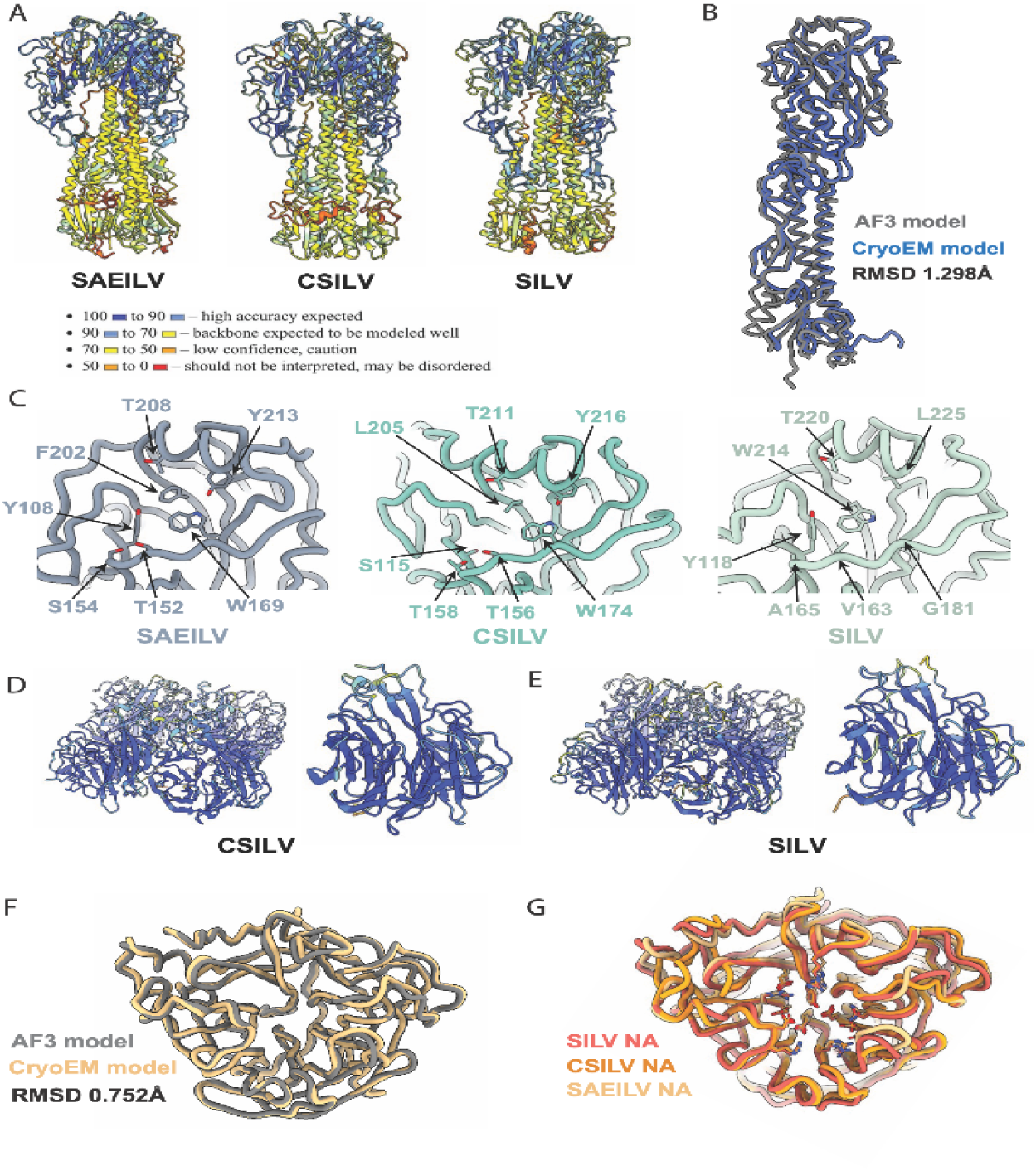
AF3 predicted HA and NA models. (**A**). AF3 predicted models of WSEIV, SAEILV, CSILV, SILV HAs, the models are colored by pLDDT scores for the confidence of accuracy. (**B**). Aligned WSEIV HA monomers from experimental data and AF3 prediction, RMSD-1.298 Å (**C**). Key residues within the RBS of SAEILV, CSILV, SILV HAs from AF3 predicted models. AF3 predicted models of CSILV (**D**) and SILV (**E**) NAs, the models are colored by pLDDT scores for the confidence of accuracy, with the same legend in (**A**). (**F**). Aligned SAELIV NA monomers from experimental data and AF3 prediction, RMSD-0.752 Å. (**H**). Aligned NA models of SAEILV (CryoEM), CSILV (AF3), and SILV (AF3) showing conserved key residues in the NA active site.

**Fig. S2.**
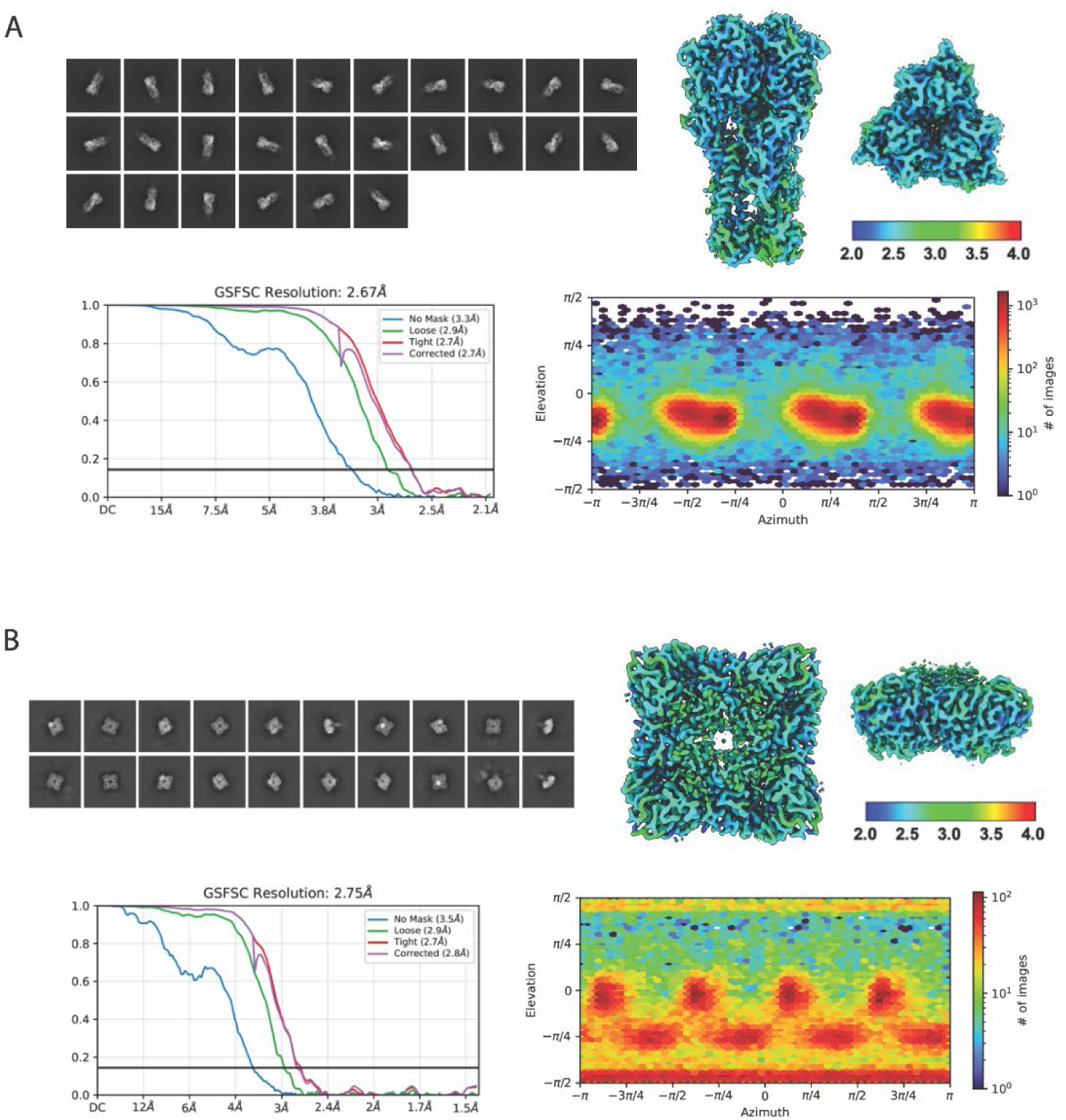
CryoEM data processing/map reconstruction. Selected 2D classes, local resolution maps, Fourier shell correlation curves, and angular sampling for WSEIV HA (**A**) and SAEILV NA (**B**).

**Fig. S3.**
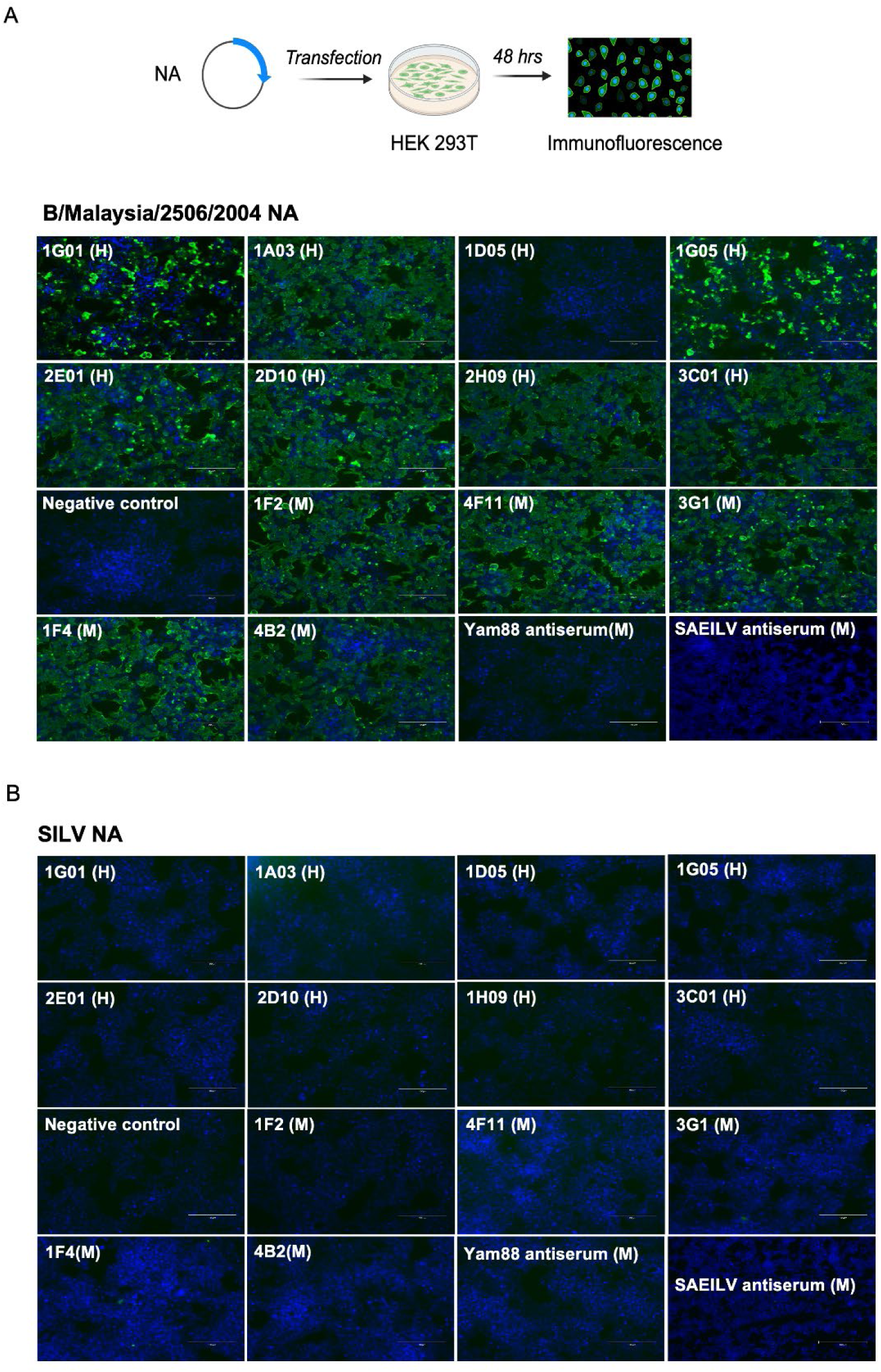
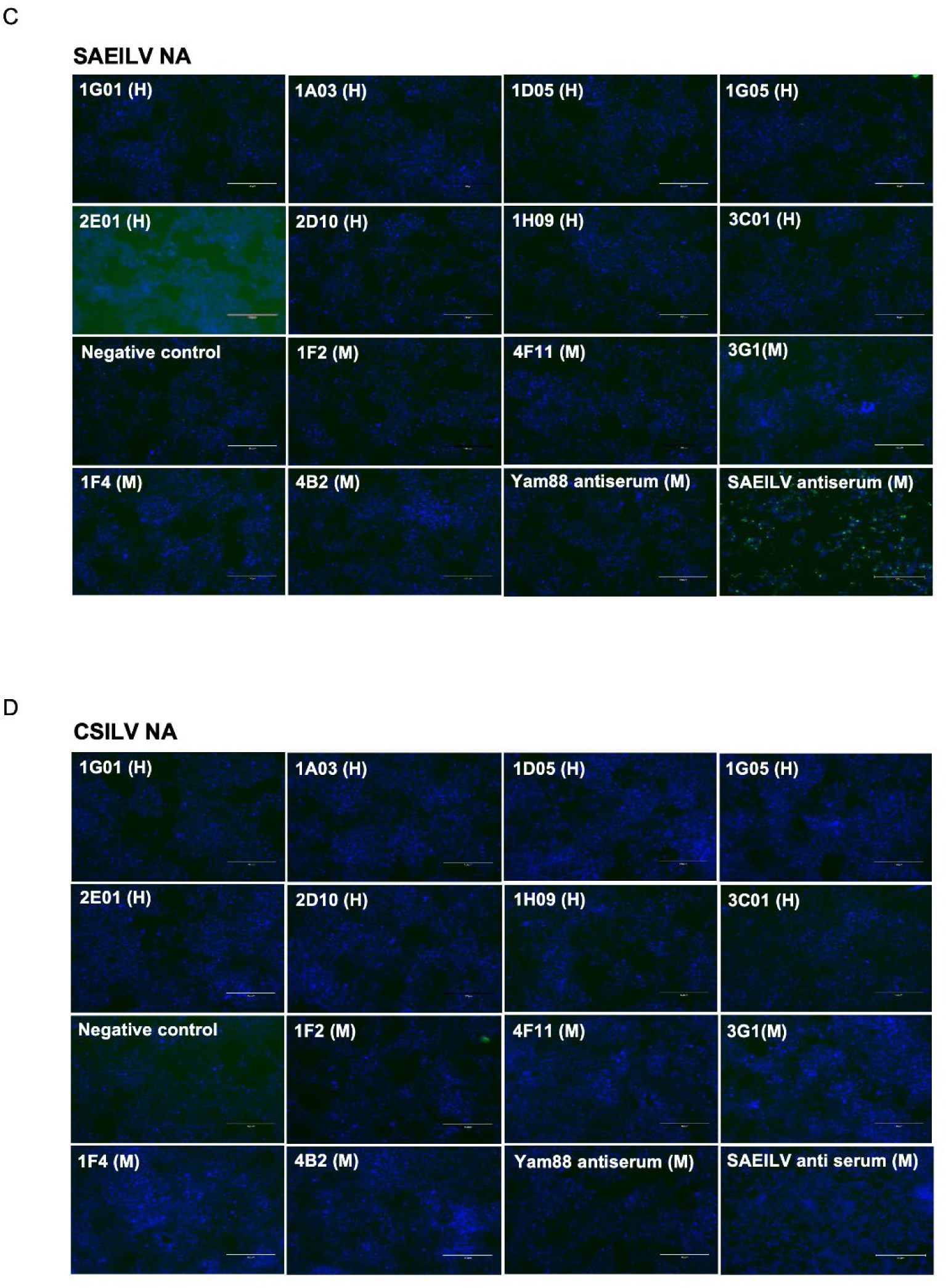
Binding of mAbs to cell expressing NAs. (**A**) Experiment setup: HEK 293T cells were co transfected with plasmid encoding full length of corresponding NA. After 48 hours cells were fixed with 3.7% PFA in PBS and incubated with human and mouse mAbs, subsequently incubated with secondary antibodies and binding was tested by immunofluorescence. Anti-influenza B virus human and mouse monoclonal antibody binding to cell surface expressed NAs of SILV(**A**), SAEILV (**B**), CSILV (**C**), and B/Malaysia/2506/2004 (**D**) by immunofluorescence.

**Table. S1.** Comparison of RBS residues of different HA subtypes and strains Table. S2 CryoEM/nsEM data collection, processing and model building statistics.

| HA | RBS residues |  |  |  |
| --- | --- | --- | --- | --- |
| H1 | 98Y | 150W | 181H | 192Y |
| H3 | 114Y | 169W | 199H | 211Y |
| H5 | 98Y | 153W | 183H | 195Y |
| B/Vic | 95F | 158W | 194H | 205Y |
| B/Yam | 95F | 158W | 193H | 204Y |
| WSEIV | 100N | 163W | 196W | 207Y |
| SAEILV | 108Y | 169W | 202F | 213Y |
| CSILV | 115S | 174W | 205L | 216Y |
| SILV | 118Y | 181G | 214W | 225L |

**Table S2.** EM data collection, processing and model building statistics.

| Map | WSEIV HA | SAEILV NA | SAEILV HA<br>(negative stain) | SILV HA<br>(negative stain) |
| --- | --- | --- | --- | --- |
| EMDB | EMD-70149 | EMD-70148 | EMD-70150 | EMD-70151 |
| **Data collection** |  |  |  |  |
| Microscope | TFS Titan Krios | TFS Glacios | TFS Talos | FEI Tecnai Spirit |
| Voltage (kV) | 300 | 200 | 200 | 120 |
| Detector | Gatan K3 | TFS Falcon 4 | FEI Ceta | FEI Eagle |
| Recording mode | Counting | Counting | Linear | Linear |
| Nominal magnification | 105,000x | 190,000x | 73,000x | 52,000x |
| Movie micrograph pixelsize (Å) | 1.045 | 0.718 | 2 | 2.063 |
| Total dose (e<sup>-</sup>/Å<sup>2</sup>) | 49.84 | 46.85 | 27 | 30 |
| Defocus range (µm) | -0.7 to -1.7 | -0.8 to -1.8 | -2 to -2.5 | -2 to -2.5 |
| **EM data processing** |  |  |  |  |
| Number of movie micrographs | 3,569 | 2,162 | 178 | 272 |
| Number of molecular projection images in map | 198,768 | 65,094 | 46,725 | 19,579 |
| Symmetry | C3 | C4 | C3 | C3 |
| Map pixel size | 1.045 | 0.718 | 2 | 2.063 |
| Map resolution (FSC 0.143; Å) | 2.67 | 2.75 | - | - |
| Map sharpening B-factor (Å<sup>2</sup>) | -86.8 | -75.4 |  |  |
| **Structure building and validation** |  |  |  |  |
| *Model composition* |  |  |  |  |
| Non-hydrogen atoms | 11,421 | 11,612 |  |  |
| Protein residues | 1,482 | 1,544 |  |  |
| Glycans | NAG:12 | NAG:4 |  |  |
| MolProbity score | 1.09 | 1.13 |  |  |
| Clashscore | 1.59 | 3.36 |  |  |
| EMRinger score | 4.46 | 5.41 |  |  |
| d FSC model (0.5; Å) | 2.9 | 2.9 |  |  |
| *RMSD from ideal* |  |  |  |  |
| Bond length (Å) | 0.006 | 0.007 |  |  |
| Bond angles (°) | 1.196 | 1.193 |  |  |
| *Ramachandran plot* |  |  |  |  |
| Favored (%) | 96.94 | 98.18 |  |  |
| Allowed (%) | 3.06 | 1.82 |  |  |
| Outliers (%) | 0.00 | 0.00 |  |  |
| Side chain rotamer outliers (%) | 0 | 0 |  |  |
| Cβ outliers (%) | 0 | 0 | | |
| PDB | 9O5W | 9O5U | - | - |

## References

1. G. J. D. S. Florian Krammer, Ron A. M. Fouchier, Malik Peiris, Katherine Kedzierska, Peter C. Doherty, Peter Palese, Megan L. Shaw, John Treanor, Robert G. Webster & Adolfo García-Sastre, Influenza. Nature Reviews Disease Primers, (2018).

2. D. M. H. Knipe, Peter, Fields Virology 6th Edition. (Lippincott Williams & Wilkins (LWW), 2013).

3. M. R. Kirsty R. Short, Josanne H. Verhagen, Debby van Riel, Eefje J.A. Schrauwen, Judith M.A. van den Brand, Benjamin Mänz,Rogier Bodewes, Sander Herfst, One health, multiple challenges: The inter-species transmission of influenza A virus. One Health, (2015).

4. N. J. Halwe et al., Bat-borne H9N2 influenza virus evades MxA restriction and exhibits efficient replication and transmission in ferrets. Nat Commun 15, 3450 (2024).

5. A. D. Osterhaus, G. F. Rimmelzwaan, B. E. Martina, T. M. Bestebroer, R. A. Fouchier, Influenza B virus in seals. Science 288, 1051–1053 (2000).

6. C. E. van de Sandt, R. Bodewes, G. F. Rimmelzwaan, R. D. de Vries, Influenza B viruses: not to be discounted. Future Microbiol 10, 1447–1465 (2015).

7. M. E. Rosu et al., Substitutions near the HA receptor binding site explain the origin and major antigenic change of the B/Victoria and B/Yamagata lineages. Proc Natl Acad Sci U S A 119, e2211616119 (2022).

8. E. C. Holmes, F. Krammer, F. D. Goodrum, Virology-The next fifty years. Cell 187, 5128–5145 (2024).

9. R. Bodewes et al., Recurring influenza B virus infections in seals. Emerg Infect Dis 19, 511–512 (2013).

10. W. T. He et al., Virome characterization of game animals in China reveals a spectrum of emerging pathogens. Cell 185, 1117–1129 e1118 (2022).

11. L. N. Measures, R. A. M. Fouchier, Antibodies against Influenza Virus Types a and B in Canadian Seals. J Wildl Dis 57, 808–819 (2021).

12. R. Bodewes et al., No serological evidence that harbour porpoises are additional hosts of influenza B viruses. PLoS One 9, e89058 (2014).

13. M. Shi et al., The evolutionary history of vertebrate RNA viruses. Nature 556, 197–202 (2018).

14. G. A. Arunkumar et al., Functionality of the putative surface glycoproteins of the Wuhan spiny eel influenza virus. Nat Commun 12, 6161 (2021).

15. R. Parry, M. Wille, O. M. H. Turnbull, J. L. Geoghegan, E. C. Holmes, Divergent Influenza-Like Viruses of Amphibians and Fish Support an Ancient Evolutionary Association. Viruses 12, (2020).

16. S. J. Gamblin et al., Hemagglutinin Structure and Activities. Cold Spring Harb Perspect Med 11, (2021).

17. F. Broszeit et al., N-Glycolylneuraminic Acid as a Receptor for Influenza A Viruses. Cell Rep 27, 3284–3294 e3286 (2019).

18. F. Krammer et al., NAction! How Can Neuraminidase-Based Immunity Contribute to Better Influenza Virus Vaccines? mBio 9, (2018).

19. Q. Wang, X. Tian, X. Chen, J. Ma, Structural basis for receptor specificity of influenza B virus hemagglutinin. Proc Natl Acad Sci U S A 104, 16874–16879 (2007).

20. F. Krammer, Emerging influenza viruses and the prospect of a universal influenza virus vaccine. Biotechnol J 10, 690–701 (2015).

21. U. Karakus et al., H19 influenza A virus exhibits species-specific MHC class II receptor usage. Cell Host Microbe 32, 1089–1102 e1010 (2024).

22. K. Ciminski, M. Schwemmle, Bat-Borne Influenza A Viruses: An Awakening. Cold Spring Harb Perspect Med 11, (2021).

23. L. Li et al., Structural and inhibitor sensitivity analysis of influenza B-like viral neuraminidases derived from Asiatic toad and spiny eel. Proc Natl Acad Sci U S A 119, e2210724119 (2022).

24. T. Velkov, The specificity of the influenza B virus hemagglutinin receptor binding pocket: what does it bind to? J Mol Recognit 26, 439–449 (2013).

25. C. Dreyfus et al., Highly conserved protective epitopes on influenza B viruses. Science 337, 1343–1348 (2012).

26. I. Margine, P. Palese, F. Krammer, Expression of functional recombinant hemagglutinin and neuraminidase proteins from the novel H7N9 influenza virus using the baculovirus expression system. J Vis Exp, e51112 (2013).

27. F. Krammer et al., A carboxy-terminal trimerization domain stabilizes conformational epitopes on the stalk domain of soluble recombinant hemagglutinin substrates. PLoS One 7, e43603 (2012).

28. E. Kirkpatrick, X. Qiu, P. C. Wilson, J. Bahl, F. Krammer, The influenza virus hemagglutinin head evolves faster than the stalk domain. Sci Rep 8, 10432 (2018).

29. G. K. Hirst, The Agglutination of Red Cells by Allantoic Fluid of Chick Embryos Infected with Influenza Virus. Science 94, 22–23 (1941).

30. T. Li et al., Chemoenzymatic Synthesis of Campylobacter jejuni Lipo-oligosaccharide Core Domains to Examine Guillain-Barre Syndrome Serum Antibody Specificities. J Am Chem Soc 142, 19611–19621 (2020).

31. W. Garten et al., Influenza virus activating host proteases: Identification, localization and inhibitors as potential therapeutics. Eur J Cell Biol 94, 375–383 (2015).

32. M. Laporte et al., Hemagglutinin Cleavability, Acid Stability, and Temperature Dependence Optimize Influenza B Virus for Replication in Human Airways. J Virol 94, (2019).

33. J. Abramson et al., Accurate structure prediction of biomolecular interactions with AlphaFold 3. Nature 630, 493–500 (2024).

34. J. U. Baenziger, D. Fiete, Structure of the complex oligosaccharides of fetuin. J Biol Chem 254, 789–795 (1979).

35. M. Dai et al., Mutation of the Second Sialic Acid-Binding Site, Resulting in Reduced Neuraminidase Activity, Preceded the Emergence of H7N9 Influenza A Virus. J Virol 91, (2017).

36. A. M. Wu et al., Differential affinities of Erythrina cristagalli lectin (ECL) toward monosaccharides and polyvalent mammalian structural units. Glycoconj J 24, 591–604 (2007).

37. V. Sharma, V. R. Srinivas, P. Adhikari, M. Vijayan, A. Surolia, Molecular basis of recognition by Gal/GalNAc specific legume lectins: influence of Glu 129 on the specificity of peanut agglutinin (PNA) towards C2-substituents of galactose. Glycobiology 8, 1007–1012 (1998).

38. D. Stadlbauer et al., Broadly protective human antibodies that target the active site of influenza virus neuraminidase. Science 366, 499–504 (2019).

39. T. J. Wohlbold et al., Broadly protective murine monoclonal antibodies against influenza B virus target highly conserved neuraminidase epitopes. Nat Microbiol 2, 1415–1424 (2017).

40. E. Kirkpatrick et al., Characterization of Novel Cross-Reactive Influenza B Virus Hemagglutinin Head Specific Antibodies That Lack Hemagglutination Inhibition Activity. J Virol 94, (2020).

41. A. Madsen et al., Human Antibodies Targeting Influenza B Virus Neuraminidase Active Site Are Broadly Protective. Immunity 53, 852–863 e857 (2020).

42. A. J. Burnham, T. Baranovich, E. A. Govorkova, Neuraminidase inhibitors for influenza B virus infection: efficacy and resistance. Antiviral Res 100, 520–534 (2013).

43. E. A. Govorkova, J. A. McCullers, Therapeutics against influenza. Curr Top Microbiol Immunol 370, 273–300 (2013).

44. F. Krammer et al., Trichoplusia ni cells (High Five) are highly efficient for the production of influenza A virus-like particles: a comparison of two insect cell lines as production platforms for influenza vaccines. Mol Biotechnol 45, 226–234 (2010).

45. K. Tamura, G. Stecher, S. Kumar, MEGA11: Molecular Evolutionary Genetics Analysis Version 11. Mol Biol Evol 38, 3022–3027 (2021).

46. T. Li et al., Chemoenzymatic Synthesis of Tri-antennary N-Glycans Terminating in Sialyl-Lewis(x) Reveals the Importance of Glycan Complexity for Influenza A Virus Receptor Binding. Chemistry 30, e202401108 (2024).

47. E. Kirkpatrick Roubidoux et al., Novel Epitopes of the Influenza Virus N1 Neuraminidase Targeted by Human Monoclonal Antibodies. J Virol 96, e0033222 (2022).

48. C. Suloway et al., Automated molecular microscopy: the new Leginon system. J Struct Biol 151, 41–60 (2005).

49. G. C. Lander et al., Appion: an integrated, database-driven pipeline to facilitate EM image processing. J Struct Biol 166, 95–102 (2009).

50. A. Punjani, J. L. Rubinstein, D. J. Fleet, M. A. Brubaker, cryoSPARC: algorithms for rapid unsupervised cryo-EM structure determination. Nat Methods 14, 290–296 (2017).

51. K. Zhang, Gctf: Real-time CTF determination and correction. J Struct Biol 193, 1–12 (2016).

52. E. F. Pettersen et al., UCSF ChimeraX: Structure visualization for researchers, educators, and developers. Protein Sci 30, 70–82 (2021).

53. P. Emsley, K. Cowtan, Coot: model-building tools for molecular graphics. Acta Crystallogr D Biol Crystallogr 60, 2126–2132 (2004).

54. D. Liebschner et al., Macromolecular structure determination using X-rays, neutrons and electrons: recent developments in Phenix. Acta Crystallogr D Struct Biol 75, 861–877 (2019).

55. P. Conway, M. D. Tyka, F. DiMaio, D. E. Konerding, D. Baker, Relaxation of backbone bond geometry improves protein energy landscape modeling. Protein Sci 23, 47–55 (2014).

